# A long intergenic non-coding RNA regulates nuclear localisation of DNA methyl transferase-1

**DOI:** 10.1101/2020.03.11.985705

**Authors:** Rhian Jones, Susanne Wijesinghe, John Halsall, Aditi Kanhere

**Affiliations:** School of Biosciences, University of Birmingham, Edgbaston, Birmingham; Institute of Inflammation and Ageing, University of Birmingham, Edgbaston, Birmingham; Institute of Genomic Sciences, University of Birmingham, Edgbaston, Birmingham; Institute of Translational Medicine, University of Liverpool, Liverpool

## Abstract

DNA methyl-transferase-1 or DNMT1 maintains DNA methylation in the genome and is important for regulating gene expression in cells. Aberrant changes in DNMT1 activity are observed in many diseases. Therefore, understanding the mechanisms behind alteration of DNMT1 activity is important. Here, we show that *CCDC26*, a nuclear long non-coding RNA frequently mutated in myeloid leukaemia, directly interacts with DNMT1. In the absence of *CCDC26* RNA, DNMT1 is mis-located in the cytoplasm. As a result, genomic DNA is significantly hypomethylated, which is accompanied by a slower cell growth rate and increased cell death. These results point to a previously unrecognised mechanism of long non-coding RNA mediated subcellular localisation of DNMT1 and regulation of DNA methylation. These observations are significant given the importance of DNMT1 in cancer and number of other diseases.

## INTRODUCTION

In the mammalian genome, DNA is often methylated at cytosines in CpG dinucleotides. DNA methylation is one of the important epigenetic modifications needed for transcriptional regulation of genes (Jaenisch and Bird 2003). This modification plays a crucial role in many vital cellular processes such as heterochromatin formation, X-chromosomal inactivation and genomic stability (Nan et al. 1998; Watt and Molloy 1988; Csankovszki, Nagy, and Jaenisch 2001). Unsurprisingly, aberrant DNA methylation is implicated in many diseases and developmental defects (Morgan, Davies, and Mc Auley 2018; Pakneshan, Tetu, and Rabbani 2004; Eden et al. 2003; Gaudet et al. 2003; Heller et al. 2016; Daskalos et al. 2009; Wen et al. 2018; Stirzaker et al. 1997; Schmelz et al. 2005; Yu et al. 2013; Klein et al. 2013; Baets et al. 2015). Regulation of DNA methylation is therefore crucial throughout mammalian existence.

In mammals, DNMT3a, DNMT3b and DNMT1 are DNA methyltransferases that are responsible for establishing and maintaining genomic methylation in cells (Lyko 2018). DNMT3a and DNMT3b are primarily involved in establishing *de novo* DNA methylation patterns in the genome. In the very early stages of development, DNA methylation is completely eradicated in primordial germ cells, resulting in an epigenetically “blank canvas”. The DNMT3s then restore DNA methylation in a non-CpG specific and ubiquitous manner (Doherty, Bartolomei, and Schultz 2002; Rasmussen and Helin 2016; Otani et al. 2009). In contrast, DNMT1 plays a more predominant role in maintaining post-replicative methylation patterns in the long-term, by preferentially binding hemi-methylated DNA, and methylating the newly synthesised daughter strand (Lyko 2018). At late S-phase of the cell cycle, DNMT1 is targeted to replication foci, a process dependent on an additional protein, ubiquitin-like with PHD and RING finger domains 1 (UHRF1) (Bostick et al. 2007; Berkyurek et al. 2014; Bronner et al. 2019). Evidence suggests that during the replication process, DNMT1 also interacts with histone modifying enzymes such as histone deacetylase HDAC2 as well as histone methyltransferases, EZH2 and G9a (Rountree, Bachman, and Baylin 2000; Esteve et al. 2006; Wang et al. 2017).

Aberrant DNA methylation is suspected to play a role in many cancers, e.g., hepatocellular carcinoma, glioblastoma, breast cancer, squamous cell lung cancer, thyroid cancer and leukaemia (Morgan, Davies, and Mc Auley 2018; Pakneshan, Tetu, and Rabbani 2004; Eden et al. 2003; Gaudet et al. 2003; Heller et al. 2016; Daskalos et al. 2009; Wen et al. 2018; Stirzaker et al. 1997; Schmelz et al. 2005; Yu et al. 2013), and DNMT mutations are also the cause of developmental diseases such as hereditary sensory and autonomic neuropathy type 1E (HSAN1E) (Klein et al. 2013; Baets et al. 2015).

In somatic cells, DNMT1 is the most abundant and most active methyl transferase. It is a predominantly nuclear protein with a N-terminal nuclear localisation signal (NLS) stretching between 177-205 amino acid residues (Alvarez-Ponce et al. 2018). The N-terminus of DNMT1 also contains domains required for its interaction with partner proteins, including DMAP1, HP1D, G9a and PCNA (Rountree, Bachman, and Baylin 2000; Esteve et al. 2006; Fuks et al. 2003; Iida et al. 2002). The central region of DNMT1 is needed for its targeting to replication foci (Leonhardt et al. 1992), whilst the C-terminus comprises the catalytic domain required for methyl-transferase activity.

Recently, a number of studies have shown that DNMT1 function is influenced by its interactions with a number of non-protein-coding or non-coding RNA transcripts (ncRNA). Non-coding RNAs like *KCNQ1OT1* (Mohammad et al. 2010), *Dali* (Chalei et al. 2014), *lincRNA-p21* (Bao et al. 2015), *PARTICLE* (O’Leary et al. 2017), *ecCEBP* (Di Ruscio et al. 2013), *DACOR1* (Somasundaram et al. 2018) and *HOXA11-AS1* (Guo et al. 2019) are shown to interact with DNMT1 and modulate its activity. Here we report an interaction between DNMT1 and a long intergenic non-coding RNA (lincRNA), *CCDC26*. *CCDC26* is transcribed from a 328-kilobase gene on chromosome 8, from a region (8q24.21) neighbouring the proto-oncogene c-MYC. The 8q24 locus has been well established as a hotbed for disease-associated mutations including cancer-associated SNPs and copy-number alterations. CCDC26 is of specific interest because it appears to be targeted by many of these mutations and variants, particularly in acute myeloid leukaemia (Radtke et al. 2009; Kuhn et al. 2012; Duployez et al. 2018; Izadifard et al. 2018). These observations suggest that *CCDC26* might play an important role in driving cancer progression. Previous studies show that *CCDC26* might be involved in regulating apoptosis and differentiation in myeloid cells (Yin et al. 2006, Hirano et al. 2008, Hirano et al. 2015). However, the exact mechanism by which *CCDC26* regulates gene expression remains elusive.

In this study, we show that *CCDC26* interacts with DNMT1 and is predominantly localised on the nuclear periphery. In the absence of CCDC26, DNMT1 is mis-localised in the cytoplasm, leading to DNA hypomethylation and apoptosis similar to that observed on inhibition of DNMT1 in myeloid leukaemia cells. As a result, we observe genome-wide changes in the expression of DNMT1 regulated genes. LincRNA mediated DNMT1 re-localisation has not been previously reported and has significant implications to DNA methylation regulation, as well as cancer and RNA biology.

## RESULTS

### LincRNA *CCDC26* is a myeloid-specific RNA expressed from second TSS

We first sought to understand the gene structure and expression pattern of CCDC26. Previous investigations (Yin et al. 2006; Hirano et al. 2015) on various leukaemia cell lines showed that *CCDC26* expression is limited to myeloid leukemia. We further examined this by measuring *CCDC26* levels in a number of additional leukaemia and non-leukaemia lines (Supplementary Figure 1A). In agreement with previous results, *CCDC26* is expressed in myeloid cells and its presence is negligible in other cell types tested. We also observed that among myeloid cells the highest level of expression is in chronic myeloid cells, K562 (Supplementary Figure 1A).

According to current gene annotations, *CCDC26* has four isoforms, which are transcribed from human chromosome 8 from two distinct transcription start sites, TSS1 and TSS2 (Figure 1A). These isoforms have alternative splicing patterns and contain combinations of six exons. Isoforms-1 (1666bp) and −2 (1649bp) are transcribed from an independent transcription start site (TSS2) and differ only by an additional 17 nucleotide sequence at the 3’ end of exon 4 in Isoform-1. Isoform-3 (1495bp), which is also transcribed from TSS2, lacks exon 4 completely. Isoform 4 (1718bp) is transcribed from TSS1, a start site that lies upstream of TSS2.

**Figure 1:**
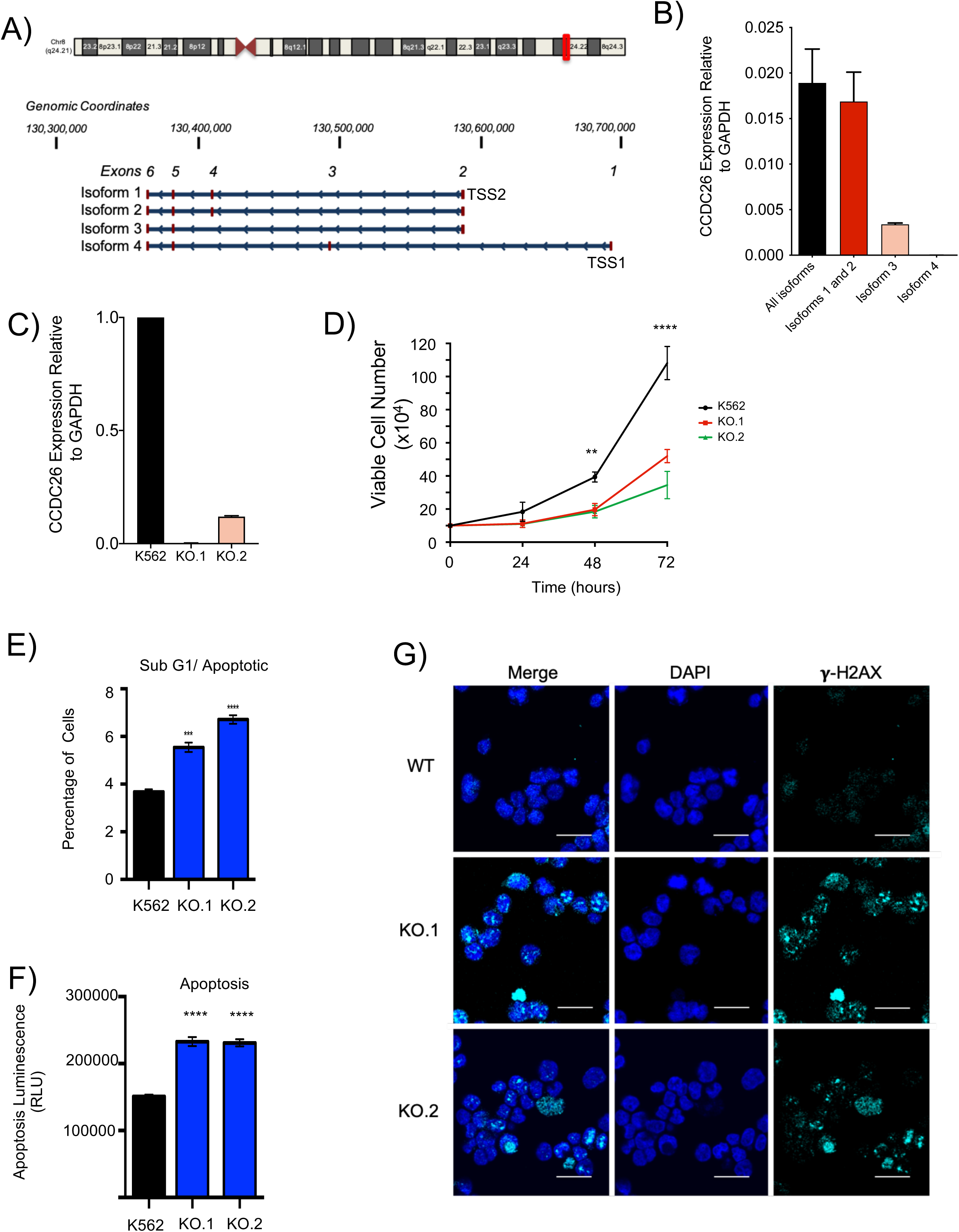
LincRNA *CCDC26* knockout results in slower growth and increased apoptosis and DNA damage in K562 cells. (A) Schematic illustrating the four currently known isoforms of *CCDC26*, adapted from the UCSC Genome Browser view of the *CCDC26* locus (NCBI RefSeq tracks, GRCh37/hg19 assembly). Isoforms are transcribed from one of two transcription start sites (TSS-1 and TSS-2) located on Chr8q24 locus. **(B)** A plot showing the RNA levels of different *CCDC26* isoforms, relative to GAPDH, in K562 cells, measured using qRT-PCR. **(C)** A plot showing expression level of *CCDC26* RNA relative to GAPDH in WT K562, KO.1 and KO.2 cells measured using qRT-PCR. *CCDC26* levels were significantly reduced in both knockouts. **(D)** Growth curve for WT K562, KO.1 and KO.2 cells showing slower growth in the latter cell lines. Values represent the mean ± standard deviation. * P<0.05; **P<0.005; ***P<0.001; ****P<0.0001 (unpaired, two-tailed *t* test, n=3). **(E)** Plot showing the percentage of apoptotic WT, KO.1 and KO.2 cells in the sub-G1 phase of the cell cycle according to propidium iodide FACS analysis. Values represent the mean ± standard deviation. * P<0.05, **P<0.005; ***P<0.001; ****P<0.0001 (unpaired, two-tailed *t* test, n=3). **(F)** The ApoTox-Glo Assay allows for detection of apoptosis in cells. The Caspase-Glo 3/7 Reagent results in cell lysis, leading to caspase cleavage and the release of the luciferase substrate amino-luciferin, causing the luciferase reaction to emit light, indicating apoptosis. Luminescence was detected and measured for WT and CCDC26 KO cells to determine the level of apoptosis in cells. Values represent the mean ± standard deviation. * P<0.05; **P<0.005; ***P<0.001; ****P<0.0001 (unpaired, two-tailed *t* test, n=5). **(G)** Confocal images demonstrating the results of anti-γ-H2AX immunofluorescence. WT, KO.1 and KO.2 cells were stained with DAPI nuclear stain (blue) and anti-γ-H2AX antibody (cyan). Increased numbers of γ-H2AX foci are present in the KO cells. Scale bar = 25um.

To understand which *CCDC26* isoforms are expressed in K562 cells, we carried out qRT-PCR measurements using isoform-specific primers (Figure 1B, Supplementary Table 1). In these cells, the four isoforms of *CCDC26* are not uniformly expressed. Isoforms starting from TSS2 i.e. isoforms −1, −2 and −3 account for more than 95% of the total *CCDC26* transcripts in the cell. Among the three isoforms starting at TSS2, Isoform-1 and Isoform-2 alone make 80% of *CCDC26* while Isoform-3 is detected at considerably lower levels, accounting for only ∼15% of *CCDC26* expression. On the other hand, Isoform-4, which is expressed from another transcription start site, TSS1, is barely detectable and represents less than 1% of total *CCDC26* (Figure 1B).

### LincRNA *CCDC26* depletion leads to DNA damage and apoptosis

To further understand the role of *CCDC26* in myeloid cells, we carried out CRISPR-Cas9 mediated knockout (KO) of this lincRNA in K562 cells. Given that ∼99% of *CCDC26* is transcribed from TSS2, we simultaneously used two small guide RNAs to mutate TSS2 (Supplementary Figure 1B). Following single cell clonal expansion, we established two cell lines; KO.1 and KO.2 which exhibit ∼99% and ∼88% reduction in *CCDC26* levels respectively (Figure 1C).

We first analysed the effect of knockout on cell growth. Interestingly, *CCDC26* depletion resulted in a significantly reduced rate of growth in both KO cell lines (Figure 1D). Cells were counted every 24hrs across a 72hr period and growth curves were subsequently plotted. Whilst wildtype (WT) cells double in number approximately every 24hrs as previously reported (Murray et al. 1993), the number of KO cells only increase two-fold approximately every 48hrs (Figure 1D).

The slow growth observed in KO cells could be because of changes in cell cycle, or because of an increased rate of cell death. Previously, *CCDC26* has been implicated in apoptosis, however its effect on cell cycle has not been investigated. To understand the reason behind slow growth rate in KO cells, we subsequently analysed cell cycle progression in WT and KOs by incorporation of Propidium Iodide and subsequent FACS analysis. This method measures the number of DNA strands to determine cell cycle progression. However, we did not observe any significant changes in any of the key cell cycle stages (Supplementary Figure 1C, D). However, cell cycle analysis showed an increased population of *CCDC26* KO cells in the sub-G1 state, which is indicative of apoptosis (Figure 1E). Cell cytotoxicity and apoptosis assays which measure caspase levels also confirmed that in comparison to WT, *CCDC26* KO cells were less viable and more apoptotic (Figure 1F). Although, the mechanism has not been studied in detail, previous publications have pointed out a role of *CCDC26* in apoptosis (Yin et al. 2006; Wang, Hui, et al. 2018; Peng and Jiang 2016; Hirano et al. 2015). The process of apoptosis is often linked to DNA damage. Hence, we explored the possibility of increased DNA damage in *CCDC26* KO cells. To verify this, we tested for increased presence of γ-H2AX (Figure 1G). Histone variant H2AX is key in the cellular response to DNA damage, having its C-terminal tail rapidly phosphorylated at a unique Serine residue following the occurrence of a DNA double-strand break (DSB). The result is the formation of γ-H2AX, which serves as an excellent DSB marker, and can be readily detected by its corresponding antibody (Rogakou et al. 1998). A visible increase in γ-H2AX foci was observed in both *CCDC26* KO lines as compared to WT cells (Figure 1G). Together, these results indicated that removal of *CCDC26* results in DNA damage, apoptosis and slow growth.

### Absence of CCDC26 leads to DNMT1 mis-localisation and DNA hypomethylation

We further sought to understand the mechanism behind *CCDC26* mediated DNA damage. We first enquired if this RNA influences genomic DNA and chromatin in any other way. Many lncRNAs are involved in regulation of chromatin modifications (Mohammad et al. 2010; Di Ruscio et al. 2013; Pandey et al. 2008; Rinn et al. 2007). We speculated that *CCDC26* might also function by regulating changes in epigenetic modifications. In order to verify this, we tested global genomic levels of multiple common histone modifications such as H3K27me3, H3K27ac, H3K9me3, H3K9ac, H4K16ac using immunoblotting (Supplementary Figure 2). However, we did not see any significant changes in any tested histone modifications in *CCDC26* KO cells. Using immunoblotting, we also tested levels of histone modifying enzymes such as EZH2, G9a and HDAC2 (Supplementary Figure 2) that are involved in catalysing these modifications. However, similar to histone modifications, these catalytic proteins, did not show any changes. In addition to histone modifications, DNA methylation is an important epigenetic modification. Importantly, DNA methylation is also shown to be regulated by various lncRNAs (Merry et al. 2015; Di Ruscio et al. 2013; Wang et al. 2015; Mohammad et al. 2010; Bao et al. 2015; Sun et al. 2016; Chalei et al. 2014; O’Leary et al. 2017; Guo et al. 2019; Gao et al. 2019). To test if DNA methylation levels have changed in the KOs, we carried out immunofluorescence measurements using anti-5-methyl Cytosine antibody (Figure 2A and Supplementary Figure 3A). Surprisingly, in KO lines, the 5-methyl Cytosine signal was considerably weaker as compared to WT cells, indicating genomic DNA was hypomethylated (Figure 2A and Supplementary Figure 3A).

**Figure 2:**
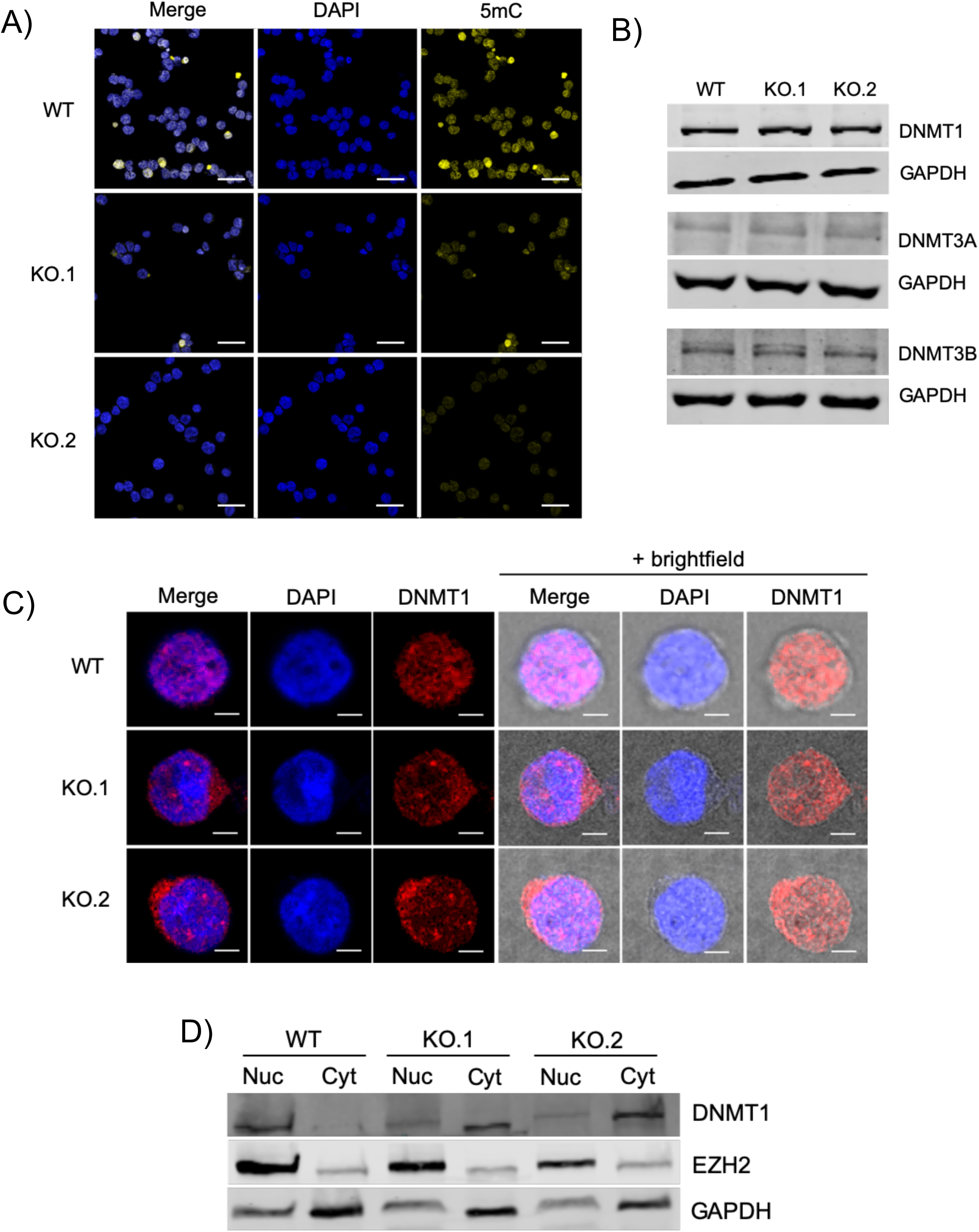
*CCDC26* knockout results in mis-localization of DNMT1 in the cytosol and global DNA hypomethylation. **(A)** Confocal images demonstrating the results of anti-5mC immunofluorescence. WT, KO.1 and KO.2 cells were stained with DAPI nuclear stain (blue) and anti-5mC antibody (yellow). Reduced levels of 5mC fluorescence are observed in both KO cell lines. Scale bar = 50um. **(B)** Total protein levels of DNMT1, DNMT3a and DNMT3b measured relative to GAPDH by immunoblotting, are unchanged in WT and *CCDC26* KO cells. **(C)** Confocal images demonstrating the results of anti-DNMT1 immunofluorescence. WT, KO.1 and KO.2 cells were stained with DAPI nuclear stain (blue) and anti-DNMT1 antibody (red). The outline of the cell membrane can be seen with the addition of the brightfield lens in the right-hand panels. DNMT1 is nuclear in WT cells and is largely cytosolic in the KO cells. Scale bar = 5um. **(D)** Immunoblotting for DNMT1 on nuclear and cytosolic protein fractions shows a shift in the subcellular localisation of DNMT1. DNMT1 is almost exclusively nuclear in the WT cells but appears both nuclear and cytosolic in *CCDC26* KO cells. EZH2 and GAPDH are used as nuclear and cytosolic markers respectively (nuc = nuclear protein fraction; cyt = cytosolic protein fraction).

A potential explanation for hypomethylation of genomic DNA in *CCDC26* KO lines could be because of changes in DNA methyltransferase levels. Immunoblotting in the knockouts, showed no significant changes in the levels of the three DNA methyltransferase proteins, DNMT3a, DNMT3b and DNMT1 (Figure 2B and Supplementary Figure 3B). Therefore, we speculated that DNA binding capacity or the enzymatic activity of a DNA methyltransferase might have changed in the absence of *CCDC26*. To detect changes in the association of DNMTs with DNA, we performed anti-DNMT immunofluorescence. We observed that in KOs, the subcellular localisation of DNMT1 is predominantly cytosolic in contrast to WT where DNMT1 is, as expected, localised in the nucleus (Figure 2C). This was supported by immunoblotting measurements using anti-DNMT1 antibody on nuclear and cytosolic fractions of KOs as compared to WT (Figure 2D). However, mis-localisation was not observed in case of DNMT3a and DNMT3b or other nuclear proteins such as HDAC2 (Supplementary Figure 3C, D).

### Genes repressed by DNMT1 are up-regulated in KO cells

We further hypothesized that if DNMT1 protein mis-localises in the cytosol, then DNMT1 would be unable to carry out its primary function of methylating genomic DNA in the nucleus. Consequently, these cells should behave similarly to cells lacking DNMT1. To confirm this, we investigated expression levels of a selection of seven genes, previously shown to be significantly impacted by DNMT1 and methylation levels in myeloid cells (Figure 3A). These were Protein Tyrosine Phosphatase Non-Receptor Type 6 (PTPN6) (Li et al. 2017; Wang et al. 2017), Cyclin Dependent Kinase Inhibitor 1A (CDKN1A) (Milutinovic et al. 2004; Schmelz et al. 2005), Cyclin Dependent Kinase Inhibitor 2B (CDKN2B) (Yu et al. 2013; Herman et al. 1996), CD9 (Kim 2012), VAV1 (Ilan and Katzav 2012; Fernandez-Zapico et al. 2005) and JUNB (Yang et al. 2003; Fiskus et al. 2009) all of which have previously demonstrated upregulation in response to DNMT1 down-regulation or DNA hypomethylation (Figure 3A). We also measured levels of IGF1, as it has previously been reported that IGF1 is repressed as a result of DNMT1 inhibition (Pastural et al. 2007). qRT-PCRs demonstrated that out of 7 genes we tested, five (PTPN6, CDKN1A, CDKN2B, CD9 and VAV1) were significantly upregulated in both KO cell lines as reported in past studies on DNMT1 KD or inhibition. JUNB, albeit not significantly, also demonstrated upregulation. As expected, IGF1 was significantly downregulated in both KO cell lines. This result demonstrates similar changes in gene expression between *CCDC26* KO cells and cells in which DNMT1 has been knocked down or inhibited. Presumably this is due to the relative functional unavailability of DNMT1 in the *CCDC26* KO cells, given its predominantly cytoplasmic localisation.

**Figure 3:**
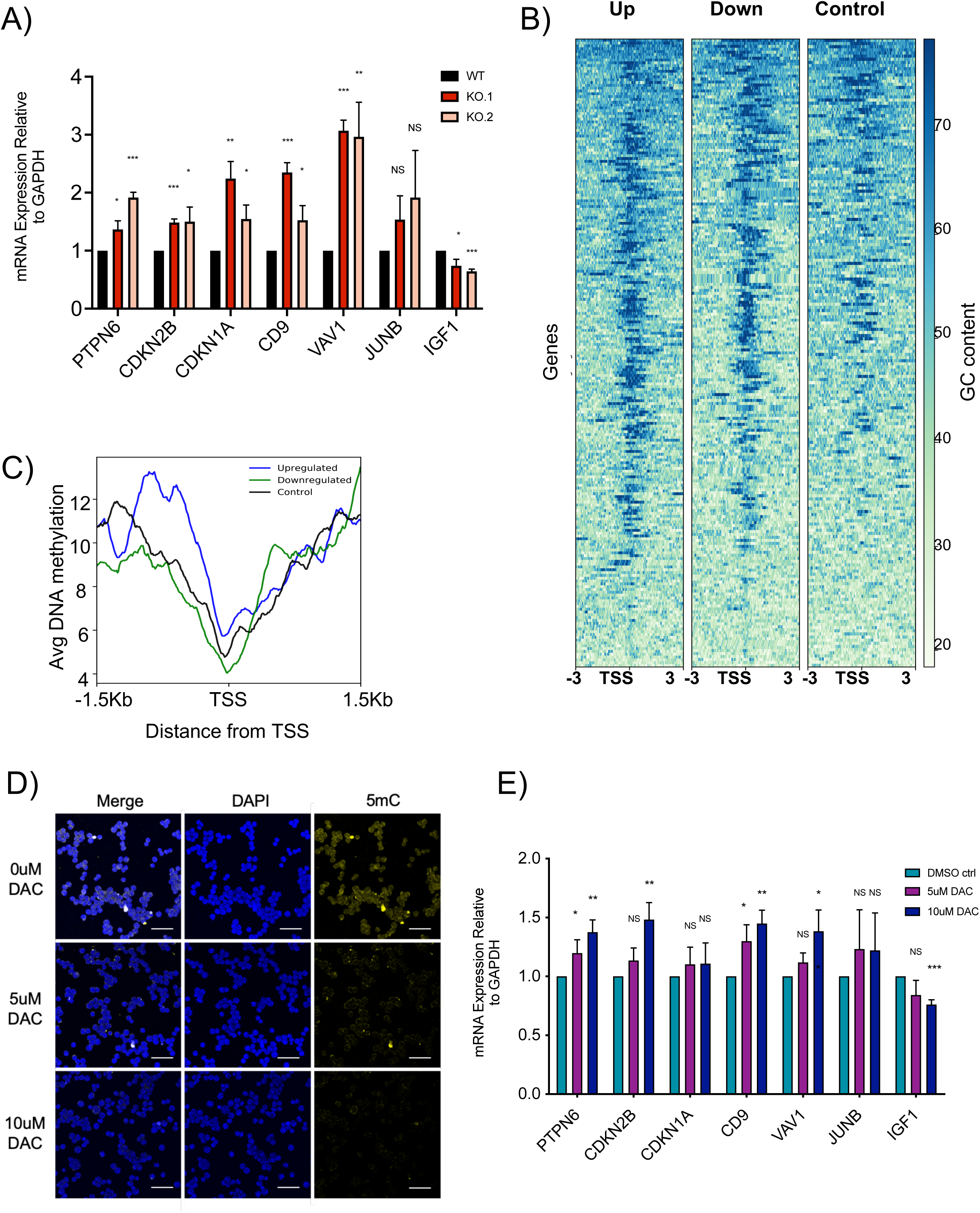
DNMT1 and methylation-regulated genes show expression changes in CCDC26 KO cells. (A) A plot showing expression levels of various genes whose expression has previously been shown to be impacted by DNMT1 depletion or DNA hypomethylation in myeloid leukemia. Levels are measured relative to GAPDH in WT K562, KO.1 and KO.2 cells by qRT-PCR Values represent the mean ± standard deviation. * P<0.05; **P<0.01; ***P<0.001; NS = Not significant (unpaired, two-tailed *t* test). **(B)** Heatmaps showing GC distribution around of Transcription Start Sites (TSS) of 2-fold downregulated and upregulated genes in KO as well as same number of randomly selected control set of genes **(C)** Metagene plots of average enrichment of DNA methylation K562 cells at 2-fold down-regulated genes (blue) as compared to up-regulated genes (green) in JARID2 KO cells and random set of genes as control (black). The plots are centered on TSS of genes and distance from TSS is indicated on the x-axes. **(D)** Confocal images demonstrating the results of anti-5mC immunofluorescence on cells treated with either 0uM, 5uM or 10uM DNMT1 inhibitor DAC. Cells were stained with DAPI nuclear stain (blue) and anti-5mC antibody (yellow). Reduced levels of 5mC fluorescence are observed in cells treated with 5uM and 10uM DAC. Scale bar = 50um. **(E)** A plot showing expression levels of various genes (as in A) whose expression have previously been shown to be impacted by DNMT1 depletion or DNA hypomethylation in myeloid leukemia. Levels are measured relative to GAPDH in cells treated with 0uM, 5uM and 10uM DNMT1 inhibitor DAC. Values represent the mean ± standard deviation. * P<0.05; **P<0.01; ***P<0.001; NS = Not significant (unpaired, two-tailed *t* test).

In order to verify this at the genomic level, using RNA-seq we measured genome-wide changes in gene expression. Our analysis showed that 287 number of genes showed more than 2-fold changes in RNA levels in both KO lines. Among these, 146 were upregulated and 141 were downregulated. It was previously shown that DNA methylation of CpG rich promoters leads to transcriptional repression (Kass, Landsberger, and Wolffe 1997; Venolia and Gartler 1983; Li, Beard, and Jaenisch 1993; Walsh, Chaillet, and Bestor 1998; Siegfried et al. 1999). If gene expression changes in *CCDC26* KO are due to mis-localisation of DNMT1, promoter methylation-mediated repression of genes should be relieved in these cells. In other words, promoters of the genes upregulated in the KOs, should be CpG-rich and should show a higher level of DNA methylation in WT cells. In order to verify this, we plotted CpG density at differentially expressed genes against a random set of genes (Figure 3B). We observed that both up- and down-regulated genes showed a much higher density of CpGs compared to the control set of genes. In addition, we utilised previously published genome-wide DNA methylation levels in K562 cells to verify if the promoters of affected genes are methylated. As suspected, genes upregulated in KO cells show a much higher level of DNA methylation in K562 cells (Figure 3C), supporting the idea that gene expression changes we see in the KOs are due to changes in DNA methylation levels imposed by mis-localisation of DNMT1 in the cytoplasm.

In order to further confirm that genes affected in the KO are regulated by DNMT1, we treated K562 cells with DNMT1 inhibitor, 5-aza-2’-deoxycytidine, also known as Decitabine (DAC). DAC is a DNMT1 inhibitor which functions by covalently trapping DNMT1 to the DNA, thereby rendering it non-functional (Stresemann and Lyko 2008). K562 WT cells were grown in the presence 0uM, 5uM and 10uM DAC for 48hrs. Treatment with both 5uM and 10uM DAC concentrations significantly reduced global 5mC levels (Figure 3D). DAC has also been shown to cause reductions of DNMT1 levels, as it is targeted for degradation after being mobilised (Ghoshal et al. 2018; Patel et al. 2010; Ghoshal et al. 2005). Western blotting for DNMT1 also demonstrated decreased protein levels (Supplementary Figure 4A).

Following confirmation of DNMT1 inhibition and global hypomethylation by DAC (Figure 3C), we next examined whether this elicited similar effects to *CCDC26* KO. Similar to the KOs, γ-H2AX immunofluorescence increased upon DNMT1 inhibition indicating an increase in DNA damage in cells treated with both 5uM and 10uM DAC for 48hrs (Supplementary Figure 4B). Significantly, qRT-PCRs for DNMT1 regulated genes showed similar patterns of expression in KO cells. Importantly, changes in gene expression were more pronounced in cells treated with 10uM DAC as compared 5uM DAC (Figure 3E). These results indicated that a DNMT1 reduction can lead to DNA hypomethylation and subsequently DNA damage.

### Apoptosis and DNA Damage is a consequence of DNMT1 mis-localisation

We next sought to further establish the sequence of events that results in DNMT1 mis-localising to the cytosol and DNA hypomethylation in *CCDC26* KO cells. It is important to examine whether cytosolic localisation of DNMT1 is a consequence of DNA hypomethylation, DNA damage and apoptosis in order to fully understand the functional mechanism of *CCDC26*.

To further confirm that DNMT1 mis-localisation is a result of *CCDC26* KO and not a consequence of DNA damage. It was critical to establish whether this type of movement of DNMT1 is not a general response to DNA damage and apoptosis. To investigate this, DNA damage was induced in WT cells using cisplatin. Cisplatin is a platinum-based drug that forms bonds with, and ultimately crosslinks, bases within and between DNA strands. This can distort the double helix, interfere with both DNA replication and transcription and consequently induce DNA damage and apoptosis (Goodsell 2006). To begin, the amount of cisplatin and treatment time required to induce DNA damage but prior to complete cell death was optimised (Supplementary Figure 5). Microscopic observations demonstrated that after 24hrs of cisplatin treatment, cells treated with 5uM or 10uM of the drug still appeared viable. DNA damage was also confirmed in the cells treated with 5uM and 10uM cisplatin by monitoring γ-H2AX foci using immunofluorescence (Figure 4A).

**Figure 4:**
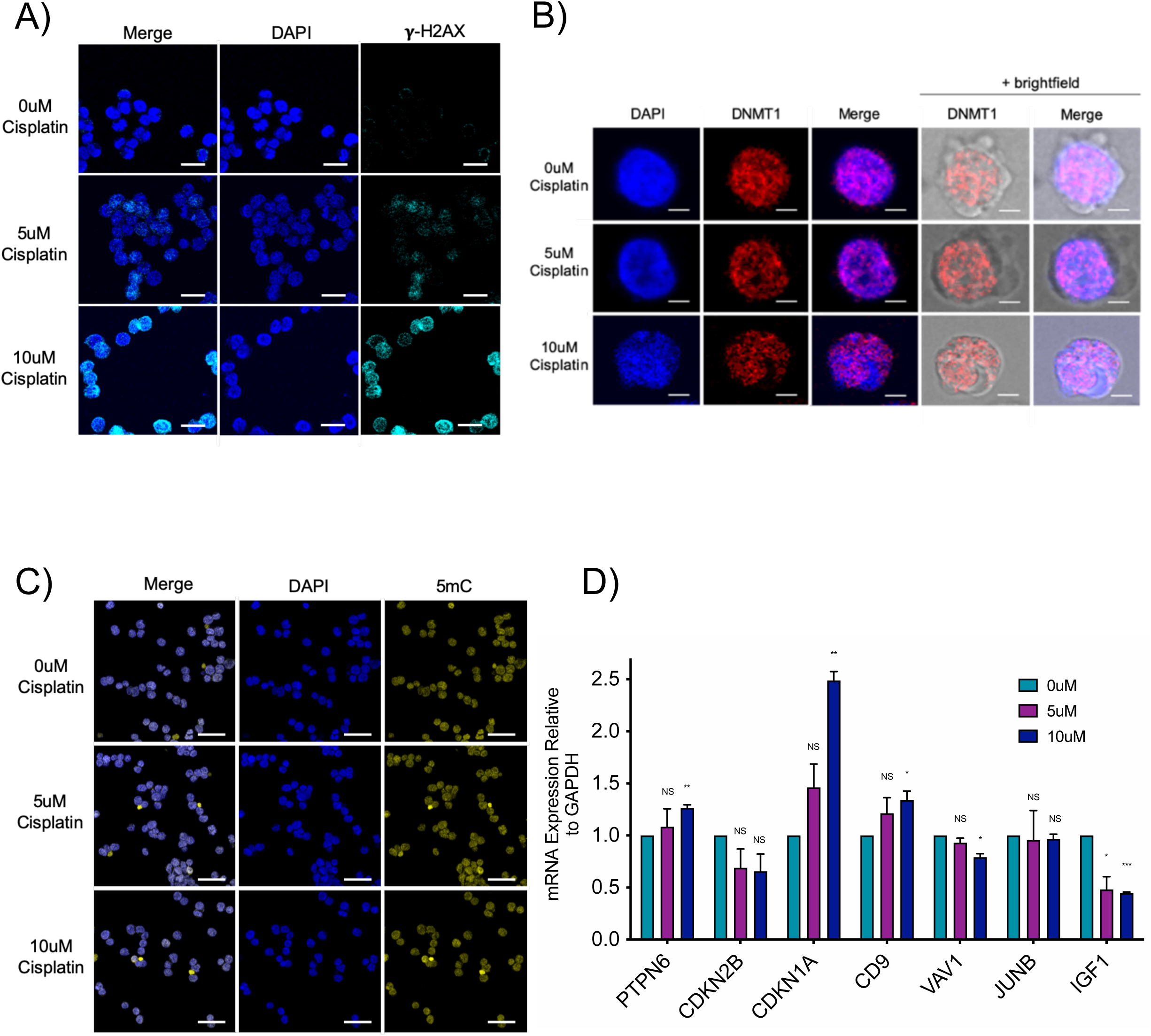
Cisplatin-induced DNA damage does not result in subcellular mis-localisation of DNMT1. (A) Confocal images demonstrating the results of anti-γ-H2AX immunofluorescence. Cells treated with 0uM, 5uM and 10uM DNA damage-inducing drug, cisplatin, were stained with DAPI nuclear stain (blue) and anti-γ-H2AX antibody (cyan). Increased numbers of γ-H2AX foci are present in the cells treated with 5uM and 10uM cisplatin. Scale bar = 25um. **(B)** Confocal images demonstrating the results of anti-DNMT1 immunofluorescence. Cells treated with 0uM, 5uM and 10uM cisplatin, were stained with DAPI nuclear stain (blue) and anti-DNMT1 antibody (red). The outline of the cell membrane can be seen with the addition of the brightfield lens in the right-hand panels. DNMT1 appears nuclear in cells treated with different cisplatin concentrations. Scale bar = 5um. **(C)** Confocal images demonstrating the results of anti-5mC immunofluorescence on cells treated with either 0uM, 5uM or 10uM cisplatin. Cells were stained with DAPI nuclear stain (blue) and anti-5mC antibody (yellow). Similar levels of 5mC fluorescence are observed in cells treated with different cisplatin concentrations. Scale bar = 50um. **(D)** A plot showing expression levels of various genes whose expression have previously been shown to be impacted by DNMT1 depletion or DNA hypomethylation in myeloid leukemia. Levels are measured relative to GAPDH in cells treated with 0uM, 5uM and 10uM cisplatin. Values represent the mean ± standard deviation. * P<0.05; **P<0.01; ***P<0.001; NS = Not significant (unpaired, two-tailed *t* test).

To assess DNMT1 localisation in DNA damage induced cells, DNMT1 immunofluorescence was performed on cisplatin-treated cells. This experiment exhibited no significant differences between 0uM control cells and the drug-treated cells. Similar to WT K562 cells, in cisplatin-treated cells, DNMT1 appeared primarily nuclear, demonstrating a diffused pattern of distribution throughout (Figure 4B). Accordingly, there was also no substantial change in the levels of 5mC immunofluorescence between control and cisplatin-treated cells (Figure 4C). This suggests that DNMT1 re-localisation is not a general consequence of DNA damage. As further confirmation, the levels of genes that are up- or down-regulated in response to *CCDC26* KO and DNMT1 inhibition were also tested. Some genes, such as IGF1, demonstrated similar changes in expression patterns in response to cisplatin, as they did in response to DNMT inhibition and *CCDC26* KO. However, in general, gene expression was either not significantly altered, or demonstrated an opposite change in expression (Figure 4D). Ultimately, we did not see identical patterns of gene expression in cisplatin-treated cells. This suggests that these expression changes are response to DNMT1 re-localisation and DNA hypomethylation but not consequence of DNA damage.

### *CCDC26* is a nuclear lincRNA and interacts with DNMT1

To further investigate the relationship between *CCDC26* and DNMT1, we investigated the possibility that *CCDC26* interacts with DNMT1 thereby localising it in the nucleus. We then endeavoured to find out the cellular localisation of this RNA. A large proportion of lncRNAs that have shown a predominantly nuclear localisation are either associated with chromatin or enriched in nuclear sub-compartments and organelles (Clemson et al. 2009; Hutchinson et al. 2007; Werner and Ruthenburg 2015). The subcellular localisation of a lncRNA can give a clue to its functional mechanism. Previous results showed that *CCDC26* might be marginally enriched in nucleus (Hirano et al. 2015). In this previous publication, all *CCDC26* isoforms were not tested for their localisation. It is possible that only selected isoforms are nuclear, thus influencing *CCDC26* function. To understand the localisation, we measured the levels of *CCDC26* isoforms using qRT-PCRs in nuclear and cytosolic RNA fractions (Figure 5A). snoRNA U105 and GAPDH levels were used as gene markers to assess the quality of nuclear and cytosolic fractions respectively (Supplementary Figure 6A). Actin was used as a housekeeping control gene against which *CCDC26* expression could be measured, given its similar levels in both the nucleus and cytosol (Supplementary Figure 6B). Consistent with previous publications (Hirano et al. 2015), all *CCDC26* isoforms were much more enriched in the nucleus as compared to cytosol (Figure 5A).

**Figure 5:**
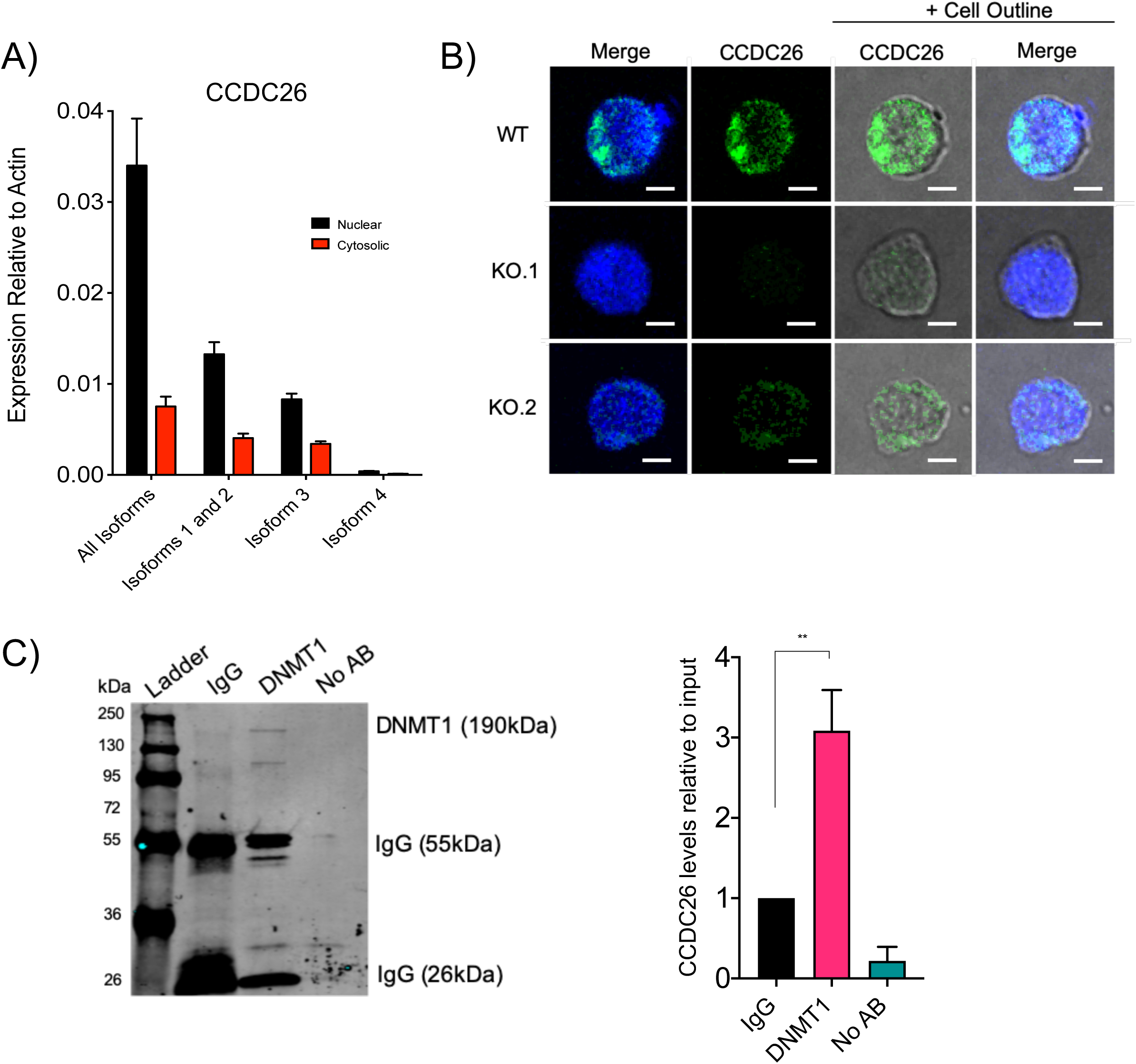
*CCDC26* is a nuclear lincRNA and interacts with DNMT1. **(A)** A plot showing the levels of different *CCDC26* isoforms, relative to Actin, in nuclear and cytosolic fractions of WT K562 cells, measured using qRT-PCR. Greater expression of all *CCDC26* isoforms was found in the nuclear fraction. **(B)** Confocal images demonstrating the results of RNA FISH using a *CCDC26*-specific fluorescent probe. WT, KO.1 and KO.2 cells were stained with DAPI nuclear stain (blue) and a *CCDC26* probe (green). The outline of the cell membrane can be seen with the addition of the brightfield lens in the right-hand panels. *CCDC26* is primarily localised in the nucleus of the cell, specifically at the nuclear periphery. Scale bar = 5um. **(C)** Protein-RNA complexes pulled down with either anti-IgG, anti-DNMT1 or no antibody, were immunoblotted with anti-DNMT1, to ensure that the DNMT1 protein was correctly pulled down for RNA immunoprecipitation. RNA pulled-down in each IP was purified, converted to cDNA and subjected to qRT-PCR with *CCDC26* primers, to determine how much *CCDC26* was pulled down relative to the input in each instance. Approximately 3X more *CCDC26* was pulled down with anti-DNMT1, compared to anti-IgG. Values represent the mean ± standard deviation. * P<0.05; **P<0.01 (unpaired, two-tailed *t* test).

To determine the location of *CCDC26* in the nucleus, we also performed fluorescence *in-situ* hybridisation (RNA-FISH) on WT and KO cells. A fluorescently labelled probe specific to exon 6 of *CCDC26* was generated and used for RNA FISH, followed by analysis using confocal microscopy. Interestingly, microscopic images (Figure 5B) clearly showed that *CCDC26* is predominantly located within the nucleus, demonstrating an enrichment at the periphery of the nucleus. The absence of fluorescent *CCDC26* signal in the KO cells indicated that the signal was specific (Figure 5B).

It is important to establish whether DNMT1 mis-localisation could be due to a direct interaction between *CCDC26* and DNMT1, or an indirect effect of *CCDC26* knockout. DNMT1 has previously been shown to bind and undergo regulation by multiple lncRNAs (Merry et al. 2015; Di Ruscio et al. 2013; Wang et al. 2015; Mohammad et al. 2010; Bao et al. 2015; Sun et al. 2016; Chalei et al. 2014; O’Leary et al. 2017; Guo et al. 2019; Gao et al. 2019). It has also been suggested that DNMT1 has higher affinity for RNA than DNA ((Merry et al. 2015; Di Ruscio et al. 2013). Nuclear localisation of CCDC26 points to a possibility that this lincRNA might also interact with DNMT1 in the nucleus. To further confirm this, we performed DNMT1 RNA immunoprecipitation (RIP) using anti-DNMT1 antibody (Figure 5C). An anti-IgG antibody was used to produce a control sample. The protein-bound-RNA pulled down with anti-DNMT1 and anti-IgG antibodies was used to perform qRT-PCRs with primers specific to *CCDC26*, and the RNA levels were measured relative to the input. Anti-DNMT1 pulled down RNA showed approximately three times more enrichment of *CCDC26* compared to the IgG control, strongly suggesting that *CCDC26* interacts with DNMT1 (Figure 5C).

A number of protein-RNA interactions have been studied using RIP or variations of this method (Di Ruscio et al. 2013; Hendrickson et al. 2016; Xu et al. 2019; Hui et al. 2019; Bai et al. 2019). At least two datasets exploring DNMT1-RNA interactions myeloid cells have been published (Di Ruscio et al. 2013; Hendrickson et al. 2016). We first adopted a bioinformatics approach where we re-mapped previously published RIP-seq data sets to the *CCDC26* locus (Supplementary Figure 6B). The first dataset that we analysed was generated using another myeloid line, HL60 (Di Ruscio et al. 2013). This data was generated by first pulling down cellular RNAs which bind to DNMT1 using anti-DNMT1 antibody and then sequencing and mapping these RNAs to the human genome. These data showed that *CCDC26* is highly enriched in DNMT1-RIP assays when compared to an IgG RIP control (Di Ruscio et al. 2013). Furthermore, analysis of another dataset, produced by a variation of the RIP method, formaldehyde-RIP-seq (fRIP-seq), also showed high enrichment of *CCDC26* in DNMT1-bound RNAs in K562 cells (Hendrickson et al. 2016), further confirming an interaction between DNMT1-*CCDC26* (Supplementary Figure 6C).

## Discussion

In the last three decades, following the development of whole-genome technologies, lncRNAs have gained importance (Hangauer, Vaughn, and McManus 2013) (Okazaki et al. 2002; Derrien et al. 2012; Iyer et al. 2015) with many examples demonstrating importance in transcription regulation.

Here we have performed functional analysis of lincRNA, *CCDC26*, in a chronic myeloid leukaemia cell line, and demonstrated its importance in regulating global DNA methylation. Our data shows that, in the absence of *CCDC26*, genome is hypomethylated which leads to increase in apoptosis and DNA damage, and cell growth inhibition. In the past, non-coding RNAs have been shown to impact DNA methylation levels through transcriptional and post-transcriptional regulation of DNMT genes (Chen et al. 2015; Wang, Guo, et al. 2018; Cheng et al. 2018; Mohammad et al. 2010; Di Ruscio et al. 2013; Merry et al. 2015). However, unlike many previous publications, DNMT expression levels were unchanged in *CCDC26* KOs, indicating that the mechanism behind the observed DNA hypomethylation is different. Strikingly, in the absence of lincRNA *CCDC26*, a large proportion of DNMT1 protein is mis-localised in the cytosol. In KO cells, the mis-localisation of DNMT1 in cytoplasm is most likely responsible for the observed hypomethylated state of the genome.

LncRNA-mediated regulation of cellular localisation has been previously reported in case of other proteins. In some instances, this has been shown to occur via a direct interaction, for example, lncRNA *TP53TG1* binds the transcription factor, YBX1, thereby preventing its nuclear trafficking (Diaz-Lagares et al. 2016). In other instances, this can also occur via an indirect effect; an interaction between lncRNA *CRYBG3* and actin for example, is sufficient to prevent translocation of Myelin and Lymphocyte protein (MAL) into the nucleus (Pei et al. 2018). Similarly, NF_k_B Interacting LncRNA (NKILA) binds and prevents phosphorylation of the inhibitory I“B subunit. This blocks its degradation, which subsequently prevents the active p65 subunit of NF_k_B from re-localising from the cytosol to the nucleus (Liu et al. 2015). However, we did not observe any changes in DNMT1 stability upon CCDC26 KO (Supplementary Figure 7). Moreover, we demonstrate an interaction between DNMT1 and *CCDC26*, suggesting that this lincRNA might play a direct role in localising DNMT1 in the nucleus.

Numerous DNMT1-interacting lncRNAs have been identified previously (Merry et al. 2015; Di Ruscio et al. 2013; Wang et al. 2015; Mohammad et al. 2010; Bao et al. 2015; Sun et al. 2016; Chalei et al. 2014; O’Leary et al. 2017; Guo et al. 2019; Gao et al. 2019), however, these largely demonstrate effects on DNA methylation at localised genes or regions, as opposed to the global effect observed here. In addition, lincRNA mediated DNMT1 localisation has not been reported before. Based on these results, we provide a model (Figure 6) suggesting that lincRNAs, in this case CCDC26, can regulate sub-cellular localisation of DNMT1 via a direct interaction in the nucleus. This provides a means of regulating global genomic DNA methylation and disruption of the lincRNA can result in hypomethylation and apoptosis.

**Figure 6:**
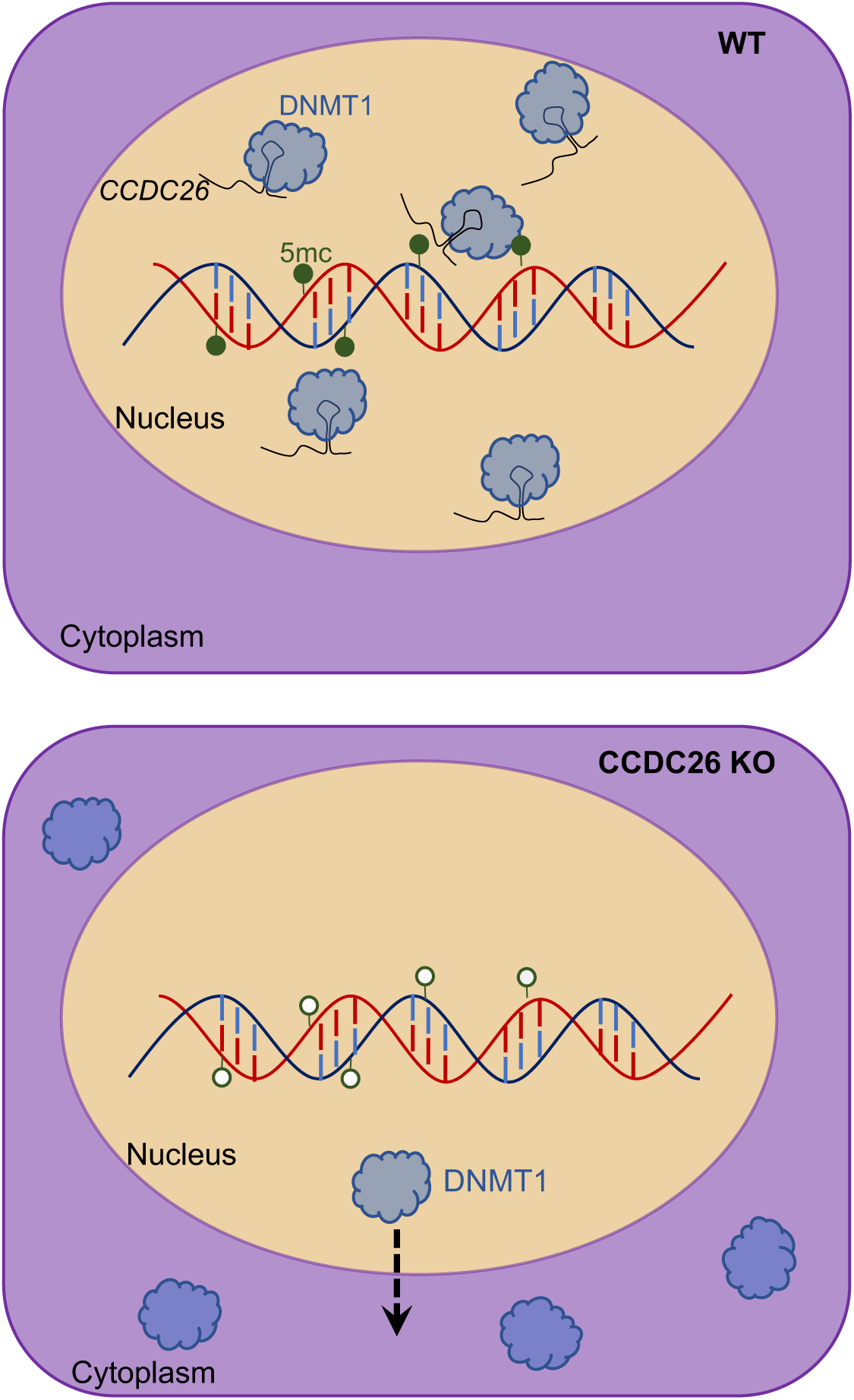
Model for *CCDC26*-mediated DNMT1 regulation. In WT K562 cells, DNMT1 interacts with *CCDC26*. DNMT1 is almost exclusively localised in the nucleus where it maintains DNA methylation patterns as cells replicate. In the absence of *CCDC26*, DNMT1 is re-localised to the cytoplasm and cells become hypomethylated.

A major question that remains in this instance, is how *CCDC26* orchestrates nuclear localisation of DNMT1, and the mechanism which causes its cytosolic mis-localisation in the absence of *CCDC26*. Independent of lincRNAs, in rare circumstances, DNMT1 localisation in the cytoplasm has been demonstrated and can provide clues regarding the mechanism behind lincRNA-mediated DNMT1 localisation. Arguably the best studied of these instances is during preimplantation in early development. An oocyte-specific form of DNMT1, DNMT1o, demonstrates a preferential localisation within the cytosol during preimplantation development. DNMT1o lacks 118 amino acids at the N-terminus compared to somatic DNMT1. It has been postulated that an alternative, extended region of the N-terminus is critical, and complex folding of this area likely plays a large part in overriding the NLS. This is required for demethylation of the embryonic genome, upon which lineage specific methylation patterns are established (Cardoso and Leonhardt 1999). In instances other than during embryonic development, cytosolic localisation of DNMT1 tends to be aberrant; for example, it has been associated with several neurological disorders including hereditary sensory and autonomic neuropathy type 1E (HSAN1E) (Baets et al. 2015), Alzheimer’s disease (Mastroeni et al. 2013) and Parkinson’s disease (Desplats et al. 2011), as well as cancer tumorigenesis (Hodge et al. 2007; Arzenani et al. 2011). The reasons behind aberrant cytosolic localisation of DNMT1 is not entirely clear. Various mechanisms including changes to post-translational modifications of DNMT1 (Hodge et al. 2007), HDAC inhibition (Arzenani et al. 2011), mutations within the RFTS domain (Baets et al. 2015) and disruption to nucleo-cytoplasmic transport systems across the nuclear membrane (Mastroeni et al. 2013) have been reported. Given that previous reports of cytosolic DNMT1 have often involved the N-terminal domain of DNMT1 (Cardoso and Leonhardt 1999; Hodge et al. 2007; Baets et al. 2015), where the NLS resides, it can be speculated that this region of the protein may be affected by the by *CCDC26*, possibly through post-translational modifications or protein folding. However, cellular and molecular mechanisms behind CCDC26-mediated DNMT1 localisation will need further investigation.

## Materials and Methods

### Cell Culture

K562 cells were maintained in an undifferentiated state in RPMI 1640 media (GIBCO, ThermoFisher Scientific), supplemented with 10% Fetal Bovine Serum (FBS) and 1% penicillin-streptomycin (10,000 U/ml), at 37°C, 5% CO2. Cells were monitored daily, passaged (split) every ∼48 hours and seeded at 2.5 x 10^5^ cells in growth media. A hemocytometer was used to observe and count cells for seeding. All cell centrifugations were performed at 1200rpm, for 5 minutes at room temperature unless stated otherwise.

To monitor cell growth, 10^5^ cells/ml were seeded into a 6 well-plate and viable cells were counted after 24, 48 and 72 hours. Trypan Blue dye exclusion was used to distinguish viable cells. Before counting, 10ul cell suspension was mixed with 10ul 0.4% Trypan Blue solution (ThermoFisher, Cat: 15250061). 10ul of this mix was then applied to a hemocytometer and cells were counted, excluding those that appeared blue.

Cells were seeded at a density of 2.5 x 10^5^ cells/ml and grown in the presence of various inhibitors and drugs. All inhibitors were prepared in a sterile tissue culture hood, and all solvents were filter sterilized before preparing stocks. Control cells were grown in the presence of the equivalent volume of solvent in which each inhibitor was dissolved (E.g. DMSO or H2O). Upon harvesting after treatment, cells were washed in PBS three times, centrifuging between washes. Cells were grown in growth media containing a final concentration of either 0uM (control), 5uM or 10uM cisplatin (Millipore, Cat: 232120) for 24 hours, before harvesting. A 10mM cisplatin stock was freshly prepared in H2O for each use. Cells were grown in growth media containing a final concentration of either 0uM (control), 5uM or 10uM 5-Aza-2’-deoxycytidine (DAC) for 48 hours, before harvesting. A 220mM DAC stock was freshly prepared in DMSO for each use.

### Propidium iodide Fluorescence Activated Cell Sorting (FACS) cell cycle analysis

For cell cycle analysis, ∼300,000 cells were centrifuged in FACS tubes and the growth media poured off. 300ul cold, cell cycle buffer (30μg propidium iodide (PI), 1% (w/v) sodium citrate, 0.1mM NaCl2, 0.1% Triton X-100) was added to the pellet, and gently vortexed, before storing at 4°C for ∼24 hours in the dark, to allow PI staining to occur. DNA content of cells was then analysed in triplicate by flow cytometry, to measure the percentage of cells at each stage of the cell cycle. This was performed using a BD FACS Calibur machine, and analysis conducted using the BD Cell Quest software (BD Biosciences).

### Cell Fixation

4 x 10^6^ cells were harvested and centrifuged at 1200rpm for 5 minutes at room temperature. Growth media was removed from the pellet, which was then washed in 1ml PBS and centrifuged again as before. The PBS was removed and the pellet was re-suspended in 50ul PBS. 1ml of a 3:1 mix of Methanol:Acetic Acid fixative was added, gently mixed and incubated at room temperature for 10 minutes exactly. This was followed immediately with centrifugation, followed by three washes with 1ml PBS. Fixed cells were stored in 3ml 70% Ethanol at 4°C at least overnight before use and stored at 4°C for no longer than 3 months.

### Immunofluorescence

Approximately 2.5×10^5^ fixed cells per slide were aliquoted into sterile microcentrifuge tubes, which were centrifuged for 5 minutes at 7000 rpm, room temperature. The supernatant was removed and the pellet was re-suspended in 200ul PBS. Samples were then spun down onto microscope slides via cyto-centrifugation for 7 minutes at 350rpm.

Cells were blocked by pipetting 50ul of 5% Bovine serum albumin (BSA) (Promega, Cat: W3841) directly onto slides, and incubating for 1 hour at room temperature, followed by a brief wash in PBS. 50ul of primary antibody (Supplementary Table 2) diluted in 5% BSA was then pipetted onto each slide and temporarily covered with a 22mm x 22mm coverslip. Slides were placed into a moist chamber and incubated overnight at 4°C. For 5mC immunofluorescence, slides were heated at 94°C for 3 minutes prior to loading primary antibody, in order to separate DNA strands to allow binding.

The following morning, coverslips were removed and slides were carefully washed three times in PBS. Approximately 80ul of the appropriate secondary antibody, diluted in PBS (Supplementary Table 3) was then pipetted onto slides, sealed temporarily with a coverslip and incubated at room temperature for 1 hour in the dark. This was followed by three PBS washes and a final single wash in H2O. After allowing slides to dry, 10ul of SlowFade® Gold anti-fade reagent (Invitrogen RNA FISH Kit, Cat. No. F32956) and 1ug/ml DAPI were added to the slide which was then covered with a coverslip and sealed with nail polish.

### RNA Fluorescence in situ Hybridisation (FISH)

A fluorescently labelled *CCDC26* RNA probe was generated using the FISH Tag RNA Multicolor Kit, Alexa Fluor dye combination (Invitrogen, Cat. No. F32956), following the manufacturers guidelines. A *CCDC26* exon 6 DNA template was generated by performing a PCR using Red Mix (Bioline, Cat. No. BIO25043) WT K562 cDNA and *CCDC26*-specific primers that incorporated a T7 RNA polymerase promoter at the 5’-end of the DNA strand to be later transcribed (Supplementary Table 1). PCR products ran on a 1% agarose gel and gel purified using the QIAquick Gel Extraction Kit (Cat. No. 28704), following the manufacturers guidelines. The subsequent *CCDC26* DNA template was subsequently used in the first in vitro transcription step of probe generation as directed by the FISH Tag RNA Multicolor Kit.

200ul fixed cells per slide were aliquoted into sterile microcentrifuge tubes, which were centrifuged for 5 minutes at 7000 rpm, room temperature. The supernatant was removed and the pellet was re-suspended in 500ul Wash Buffer A (5ml 20X nuclease free saline-sodium citrate (SSC), 5ml deionized formamide, 40ml nuclease-free H_2_O). Cells were centrifuged as before and the pellet resuspended in 100ul Hybridisation Buffer (1g dextran sulphate, 1ml deionized formamide, 1ml 20X nuclease-free SSC, 8ml nuclease-free H_2_O) containing 1ug/ml of probe. This was mixed well by pipetting and incubated overnight, in the dark at 37°C. The following morning, the cells were centrifuged and washed with 500ul Wash Buffer A. Following another centrifugation, the pellet was resuspended in 500ul Wash Buffer A and incubated in the dark, at 37°C for 30 minutes. The cells were centrifuged, and the buffer removed. Cells were then resuspended in 200ul PBS and spun down onto microscope slides via cyto-centrifugation for 7 minutes at 350rpm. 10ul of SlowFade® Gold anti-fade reagent (Invitrogen RNA FISH Kit, Cat. No. F32956) and 1ug/ml DAPI were added to the slide which was then covered with a coverslip and sealed with nail polish.

### Confocal Microscopy

Slides were imaged using a Leica TCS SP8 Confocal microscope, using either 20X, 40X or 63X objectives. The brightfield microscope setting was used to visualise cell membrane boundaries. FIJI image analysis software was used to visualise and analyse confocal images.

### RNA Extraction

An RNase-free environment was maintained for all RNA work by treatment of equipment and work surfaces with RNase *Zap*, RNase decontamination solution (ThermoFisher Scientific, Cat. No. AM9780).

All RNA was extracted using the QIAGEN RNeasy Mini Kit (Cat: 74106), following manufacturers guidelines. RNA was eluted in 30ul RNase-free H_2_O and concentrations were determined using a NanoDrop ND-100 spectrophotometer (Thermo Scientific). RNA quality was determined by running ∼200-300ng on a 1% agarose gel.

### Nuclear and Cytosolic RNA Extraction

For extraction of nuclear and cytosolic RNA fractions, approximately 10^7^ cells were harvested by centrifuging at 1200rpm at room temperature for 5 minutes, washing in 1ml PBS and centrifuging again. As much supernatant as possible was removed from the cell pellet, which was then re-suspended in 1ml of cold (4°C) Buffer RLN (50mM Tris-HCl (pH.8), 1.5mM MgCl2, 140mM NaCl, 0.5% NP-40, 100U/ml RNase inhibitor) and incubated on ice for 5 minutes. 250ul of the mix was then pipetted into four eppendorfs labelled “nuclear”. These were centrifuged for 2 minutes at 3700rpm, 4°C. Approximately 500ul of supernatant collected from all four eppendorfs was transferred into a fresh eppendorf, labelled “Cytosolic”, taking extra care not to disturb the nuclear pellets. Each of the four nuclear pellets were washed twice with 100ul Buffer RLN, centrifuging for 2 minutes at 3200rpm, 4°C between washes. 200ul and 600ul Buffer RLT (containing 1% β-mercaptoethanol, QIAGEN RNeasy Mini Kit) was added to nuclear and cytosolic fractions respectively and vortexed vigorously. The nuclear pellets were further homogenized using a 1ml syringe and needle (21G x 1.5” – Nr.2., 0.8mm x 40mm). The fractions were then centrifuged for 3 minutes at maximum speed to remove cell debris, and the supernatants were removed and mixed with 1X volume 70% ethanol. The extraction was continued using the QIAGEN RNeasy Mini Kit, following manufacturers guidelines.

All freshly extracted RNA was treated with DNase I using the Sigma, amplification grade DNase I kit (Cat: AMPD1), following the kit guidelines, and converted into cDNA using the Bioline-Tetro cDNA Synthesis Kit (Cat. No. BIO-65042) following the manufacturers guidelines.

### Quantitative Real-Time Polymerase Chain Reaction (qRT-PCR)

qRT-PCRs were performed in 96-well plates. 20ng of cDNA made up to 6.8ul with dH2O was added to each well (in triplicate) with 1.6ul forward primer (5uM), 1.6ul reverse primer (5uM) and 10ul SensiFAST SYBR Hi-ROX mix (Bioline, Cat: BIO92020). The plate was then loaded into an Agilent AriaMax real time PCR machine (Agilent Technologies) and the qRT-PCR was performed (1 cycle at 95.0°C for 2 minutes, followed by 40 cycles at 95.0°C for 5 seconds, 60.0°C for 10 seconds and 72.0°C for 10 seconds).

The average cycle threshold (Ct) values from each technical triplicate were calculated and used to determine expression levels of the gene of interest relative to the housekeeping gene, GAPDH using the formulae below: ΔCt = Ct (gene of interest) – Ct (housekeeping gene) ΔΔCt = ΔCt (sample group) - ΔCt (control group) Relative Expression = 2^-ΔΔCt^ After calculating gene expression relative to the housekeeping gene, GAPDH, unless stated otherwise, values for three biological replicates were averaged and unpaired, two-tailed, parametric t-tests were performed to calculate the significance of any differences between control and treated samples. *P* values of <0.05 were considered as statistically significant.

### Total Protein Extraction

Total protein was extracted from cells by first harvesting ∼2 x 10^6^ cells by centrifuging at 1200rpm for 5 minutes. Cell pellets were washed with 1ml PBS and centrifuged again as before. As much PBS as possible was removed from the pellets before re-suspending in ∼80ul lysis buffer (20mM Tris, pH 7.5, 150mM NaCl, 10mM EDTA, 0.5% deoxycholic acid, 0.5% Triton X-100). 1 protease inhibitor tablet (Roche, Cat. 04693159001) was added to 10ml lysis buffer immediately before use. Samples were then homogenized using a 1ml syringe and needle (21G x 1.5” – Nr.2., 0.8mm x 40mm) and incubated on ice for 30 minutes before centrifuging at 14,000rpm for 20 minutes at 4°C. The resulting supernatant was pipetted into a fresh, sterile eppendorf and stored at −20°C.

### Nuclear and Cytosolic Protein Extraction

Approximately 1.5 x 10^7^ cells were harvested by centrifuging at 1200rpm for 5 minutes at room temperature and washed in PBS. As much supernatant was removed from the pellet as possible, which was then re-suspended in 150ul cold Buffer RLN and incubated on ice for 10 minutes. One protease inhibitor tablet (Roche, Cat. 04693159001) was added to 10ml Buffer RLN immediately before use. The mix was centrifuged at 3700 rpm for 5 minutes at 4°C and 100ul supernatant was transferred to a new eppendorf (cytosolic fraction). The remainder of the supernatant was discarded and the pellet was washed twice with 100ul Buffer RLN, centrifuging between washes as before. The resulting pellet was re-suspended in ∼80ul lysis buffer. One protease inhibitor tablet (Roche, Cat. 04693159001) was added to 10ml lysis buffer immediately before use. Samples were then homogenized using a 1ml syringe and needle (21G x 1.5” – Nr.2., 0.8mm x 40mm) to lyse nuclei, and incubated on ice for 30 minutes before centrifuging at 14,000rpm for 20 minutes at 4°C. The resulting supernatant was pipetted into a fresh, sterile eppendorf (nuclear fraction) and stored at −20°C.

### Histone Protein Extraction

10^7^ cells were harvested for histone extraction. Cells were first lysed in 500ul histone extraction buffer (PBS containing 0.5% Triton X-100, 2mM phenylmethylsulfonyl fluoride, 0.02% sodium azide) for 1 minute on ice, followed by centrifuging at 8000rpm for 10 minutes at 4°C. The resulting pellet was washed in 250ul histone extraction buffer and centrifuged as before. The pellet was then resuspended in 0.4M HCl and incubated overnight at 4°C for acid extraction of histones. The following morning, samples were centrifuged as before and the supernatant (containing histone protein) was saved. The HCl was neutralized by adding 0.1X volumes of 2M NaOH. The histone protein was then used in SDS-PAGE (2.7.6). All histone extractions were performed in collaboration with Dr John Halsall (Institute of Cancer and Genomic Sciences, University of Birmingham) (Halsall et al. 2015)

### Bradford Assay Protein Quantification

To ensure equal loading of samples in SDS-PAGE, protein concentrations were determined via Bradford assay. Standard samples (5ul of known BSA protein concentration in serial dilution ranging from 0-2mg/ml) and 5ul of experimental protein samples were loaded in duplicate into a 96-well plate along with 250ul Bradford reagent (Sigma, Cat: B6916). Absorbance was measured at 570nm using a Tecan infinite 5200 pro plate reader and iControl^TM^ Microplate Reader Software. A standard curve was plotted from the standard samples of known protein concentration (conc. vs absorbance). The protein concentration of the experimental samples was determined by subtracting the absorbance from a blank control absorbance value, and then dividing by the slope of the standard curve.

### Immunoblotting

For SDS-PAGE, 10% polyacrylamide SDS resolving gels (6.66ml 30% acrylamide, 5ml 1.5M Tris pH8.8, 200ul 10% SDS, 200ul 10% APS, 20ul TEMED and 8ml H_2_O) were prepared with a 4% stack (1.7ml 30% acrylamide, 2.5ml 0.5M Tris pH6.8, 100ul 10% SDS, 100ul 10% APS, 20ul TEMED and 5.55ml H_2_O) and stored in moist wrapping at 4°C for no more than 1 week before use.

Before loading onto gels, extracted protein samples were mixed with 5X SDS loading dye (200mM Tris-HCL pH6.8, 40% glycerol, 4% SDS, 0.4% bromophenol blue, 200mM β-mercaptoethanol) and heated at 70°C for 10 minutes. In addition to protein samples, 3ul of protein ladder (PageRuler^TM^ Plus Prestained Protein Ladder, ThermoFisher Scientific, Cat: 26619) was loaded onto gels as a molecular weight marker for use as a size standard reference. Gels were then run for approximately 90-120 minutes at 120V in cold 1X Running Buffer (100ml 10X running buffer, 5ml 20% SDS, 895ml dH2O) (10X running buffer: 60g Tris, 288g glycine, up to 2L dH_2_O).

After running, a semi-dry transfer was performed onto nitrocellulose membranes using the Trans-Blot Turbo Transfer System (BIO-RAD, Cat. 1704271) and 1X Transfer Buffer (200ml BIO-RAD TransBlot Turbo 5 x Transfer Buffer (Cat. 1704271), 200ml 100% Ethanol, 600ml nanopure H2O). Membranes were then blocked in 5% skimmed milk in TBS for 1 hour before incubating in primary antibody overnight at 4°C, with agitation. All primary antibodies (Supplementary Table 2) were prepared in 5% BSA. The following morning, the membranes were washed three times in TBS-T for 5 minutes and incubated in the appropriate secondary antibody (Supplementary Table 3), diluted in blocking solution (5% skimmed milk in TBS-T) in the dark for 1 hour, at room temperature, with agitation. The antibody was removed and the membranes were washed twice in TBS-T for 5 minutes, and once in TBS for 5 minutes for a final time, before storing in TBS at 4°C.

Membranes were scanned and protein bands were detected using the Odyssey infrared detection system (LI-COR Biosciences). Images were then analysed and the protein bands were quantified using Image Studio Lite software.

### DNMT1 RNA Immunoprecipitation (RIP)

2 x 10^7^ cells were harvested per RIP. Cell pellets were washed in PBS three times and resuspended in 500ul ice-cold Buffer RLN (containing 100U/ml RNase inhibitor and with 1 protease inhibitor tablet added to 10ml RLN). This was incubated on ice for 10 minutes before centrifuging at 3700rpm, for 5 minutes at 4°C. Nuclear pellets were washed with 100ul Buffer RLN, followed by resuspending in 500ul freshly prepared RIP Buffer (25mM Tris, pH 7.4, 5mM EDTA, 150mM KCl, 0.5mM DTT, 0.5% NP40 Igepal, 100U/ml RNase inhibitor) with added protease inhibitor. This mix was incubated on ice for 3 hours with frequent, gentle agitation. Following the incubation, nuclei were homogenized using a 1ml syringe and needle (21G x 1.5” – Nr.2., 0.8mm x 40mm), then centrifuged at maximum speed for 10 minutes at 4°C to pellet nuclear debris. The supernatant was kept and transferred to an RNase-free eppendorf, to which 6ug of either DNMT1 of IgG (control) antibody was added (Supplementary Table 2). This mix was incubated overnight at 4°C with gentle rotation.

The following day, 50ul (1.5mg) magnetic beads (Dynabeads Protein G Immunoprecipitation kit, Invitrogen, Cat: 10007D) were prepared per RIP. Similar to 2.8, the beads were placed into an eppendorf on a magnetic separator and the supernatant was removed. This was followed by two washes with 100ul RIP buffer. The overnight antibody-protein-RNA suspension was added to the beads and incubated for 1 hour with gentle rotation at 4°C. Following this incubation, the beads-antibody-protein-RNA were placed on the magnet, the supernatant was removed and the complex was washed three times with 500ul ice-cold RIP buffer and once with RNase-free PBS. At this point, 5% of the bead slurry was collected to be used in SDS-PAGE analysis to confirm DNMT1 pull-down. The remaining beads-antibody-protein-RNA was resuspended in 100ul RIP buffer with 50ug proteinase K and 0.1% SDS.

This was incubated at 55°C for 45 minutes to detach the protein-RNA complexes from the beads. The eppendorf was then placed onto the magnet and the supernatant was transferred to a new eppendorf. The beads were discarded.

To purify the RNA, 1X volume of phenol-chloroform-isoamyl alcohol was added to the supernatant and vortexed thoroughly. This was phase separated by centrifuging at 14,000rpm for 10 minutes at 4°C. The aqueous phase (containing the RNA) was carefully collected and placed into a fresh eppendorf. Any remaining aqueous phase was further extracted by the addition of 150ul back extraction buffer (10mM Tris, pH8, 1mM EDTA, 100mM NaCl, 0.25% SDS), followed by vortexing and centrifugation as before. Any remaining aqueous phase was collected and added to the previous collection.

The RNA was further purified by ethanol precipitation. 0.1X volumes of 3M sodium acetate (pH 5.2), 2.2X volumes of 100% ice cold ethanol and 1ul glycogen was added to the RNA extract and incubated at −20°C overnight. The following morning, the mix was centrifuged at 10,000rpm for 20 minutes at 4°C and the supernatant carefully discarded. 500ul 70% ice cold ethanol was added and the mix centrifuged as before. The ethanol was carefully removed and the tubes were left open to allow any residual ethanol to evaporate. The purified RNA was then finally dissolved in 20ul RNase-free H2O. This RNA was then DNase-treated and converted into cDNA, and qRT-PCRs were performed. Results were analysed using the calculations below to determine DNMT1 enrichment for *CCDC26* compared to the IgG control. DNMT1 RIP ΔCt = Ct DNMT1_RIP_CCDC26 – Ct INPUT_RNA_CCDC26 IgG RIP ΔCt = Ct IgG_RIP_CCDC26 – Ct INPUT_RNA_CCDC26ΔΔCt = DNMT1 RIP ΔCt – IgG RIP ΔCt Relative Expression = 2^-ΔΔCt^

### RNA-sequencing and metagene analyses

RNA-sequencing was carried out on RNA extracted from wild-type K562 cells and CCDC26 KO K562 cells. The sequencing was carried out on three biological triplicates. RNA-sequencing libraries were prepared using TrueSeq method. All the libraries were paired-end sequenced on an Illumina HiSeq 2500 machine (University of Birmingham). Sequences were quality filtered and trimmed using cutadapt. The reads were mapped using TOPHAT package (Trapnell, Pachter, and Salzberg 2009) against human genome (hg19). Differential analysis was done using cuffdiff programme. Differential genes were identified using false discovery cut-off of 1 x 10^-5^ and used for further analysis. The sequencing data is deposited GEO database (accession no. GSE105029) Metagene plots and heatmap were generated using DeepTools package(Ramirez et al. 2014). DNA methylation metagenes were generated using K562 MeDIP data deposited in ENCODE database (GEO: GSM2424238). The GC heatmap were created using previously calculated GC content track for human genome available from UCSC genome browser (http://hgdownload.cse.ucsc.edu/goldenPath/hg19/gc5Base/).

## Acknowledgements

We thank Dr J Woolley, Dr M Winch and Dr E Petermann for useful discussions. We are also grateful to Dr Alessandro Di Maio for help with microscopy data analysis. During the course of study, AK was funded by SSfH fellowship from University of Birmingham. RJ is supported by BBSRC MIBTP fellowship. SW was supported by MRC and University of Birmingham.

## Author Contributions

AK conceived the study. RH and AK wrote the manuscript. RH, SW, JH and AK designed and carried out the experiments.

---

**Supplementary Table 1.**
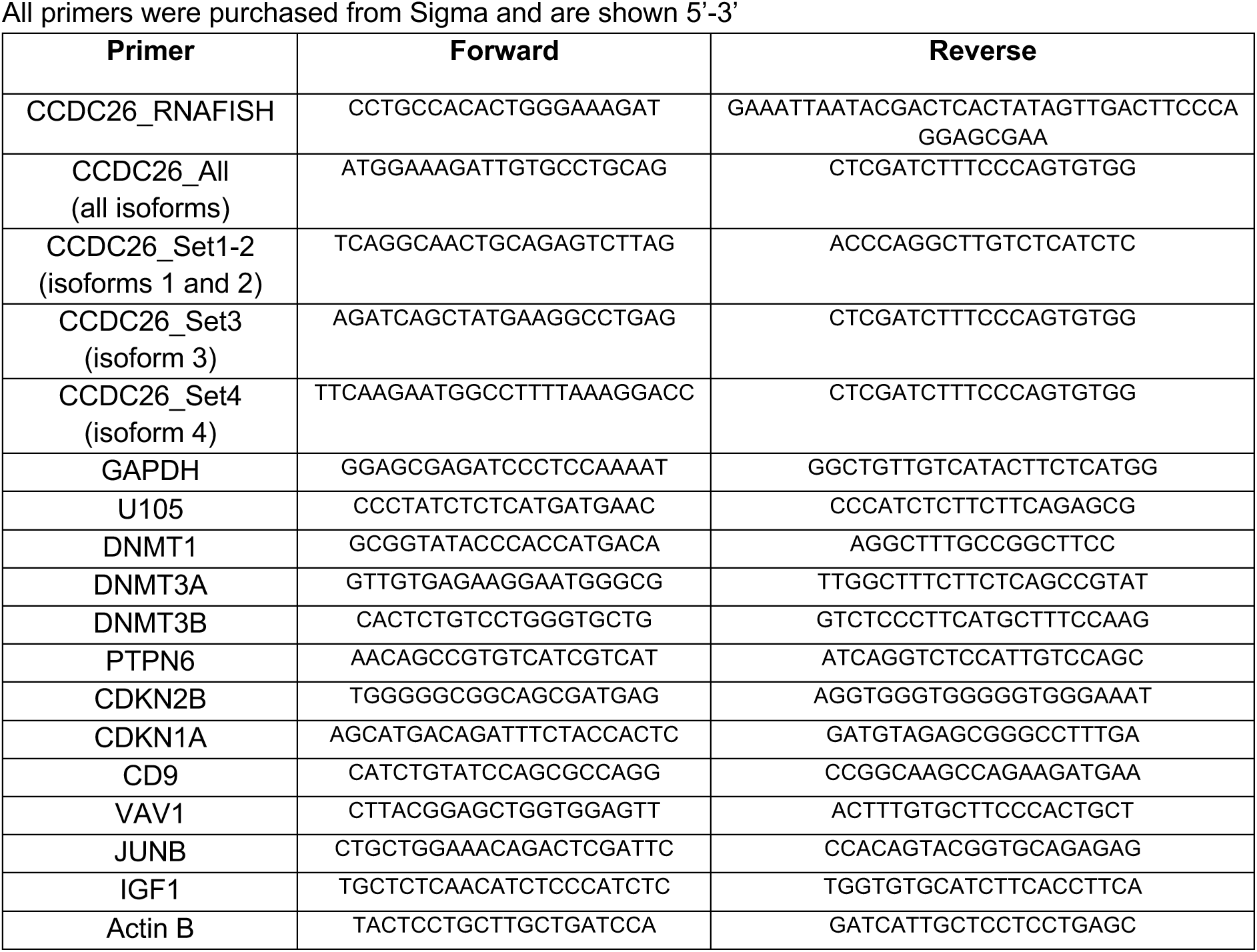
Primers All primers were purchased from Sigma and are shown 5’-3’

**Supplementary Table 2.**
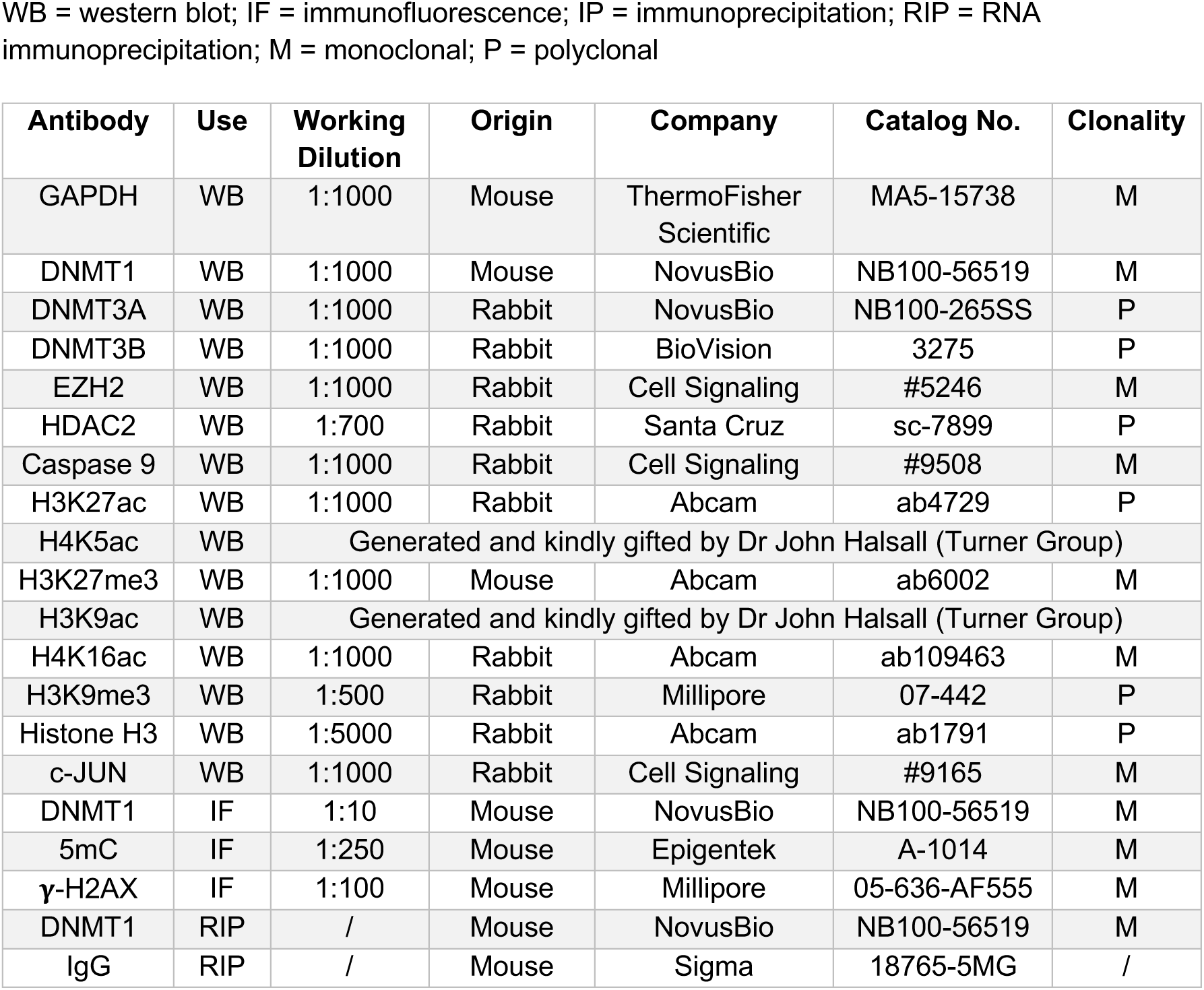
Primary Antibodies WB = western blot; IF = immunofluorescence; IP = immunoprecipitation; RIP = RNA immunoprecipitation; M = monoclonal; P = polyclonal

**Supplementary Table 3.** Secondary Antibodies WB = Western Blot; IF = Immunofluorescence

| **Antibody** | **Use** | **Working Dilution** | **Company** | **Catalog No.** |
| --- | --- | --- | --- | --- |
| **Anti-rabbit IgG (H+L)**<br>**(DyLight™ 800 4X PEG**<br>**Conjugate)** | WB | 1:10,000 | Cell Signalling | 5151 |
| **Anti-mouse IgG (H+L)**<br>**(DyLight™ 680 Conjugate)** | WB | 1:10,000 | Cell Signalling | 5470 |
| **Alexa Fluor 488 donkey anti-**<br>**rabbit IgG (H+L)** | IF | 1:500 | Invitrogen | A21206 |
| **Alexa Fluor 633 goat anti-**<br>**mouse IgG (H+L)** | IF | 1:500 | Invitrogen | 1010093 |

