## Supplementary figures for "A long intergenic non-coding RNA regulates nuclear localisation of DNA methyl transferase-1"

### SUPPLEMENTARY FIGURE -1

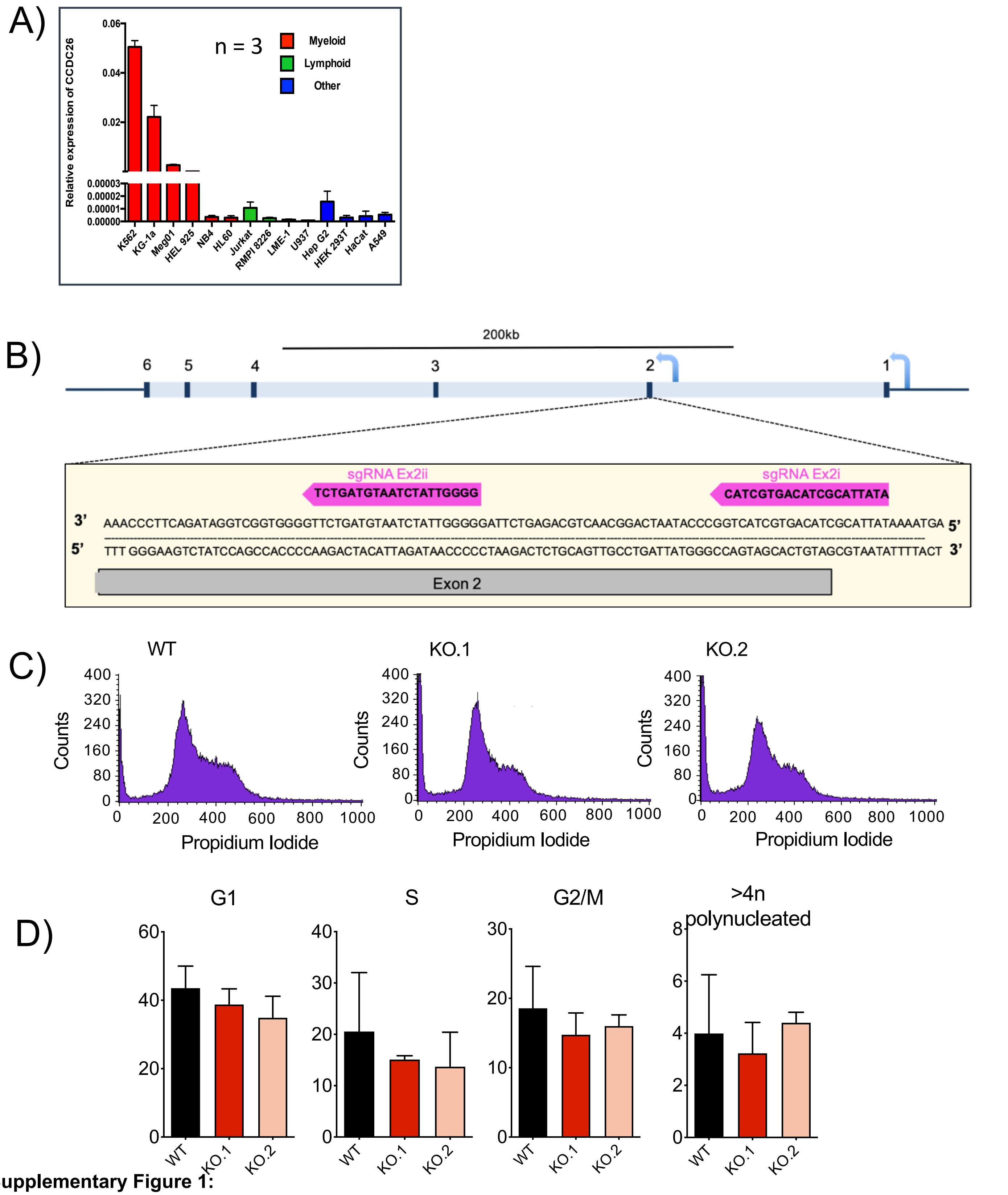


**Supplementary Figure 1:**

*(A)* *CCDC26* expression in 14 different cell-lines of diverse origins. Expression was measured using qRT-PCR assays and normalised against expression levels of Actin.

(B) Schematic of the *CCDC26* gene showing that TSS-2 was targeted with two sgRNAs simultaneously using CRISPR Cas9 technology to establish *CCDC26* KO cell lines.

(C) Histograms showing distribution of WT and KO cells in different cell cycle stages after propidium iodide staining and FACS analysis.

(D) Plots showing the percentage of WT, KO.1 and KO.2 cells in each stage of the cell cycle following propidium iodide staining and FACS analysis. Values represent the mean ± standard deviation (n=3) (unpaired, two-tailed *t* test).

### SUPPLEMENTARY FIGURE -2

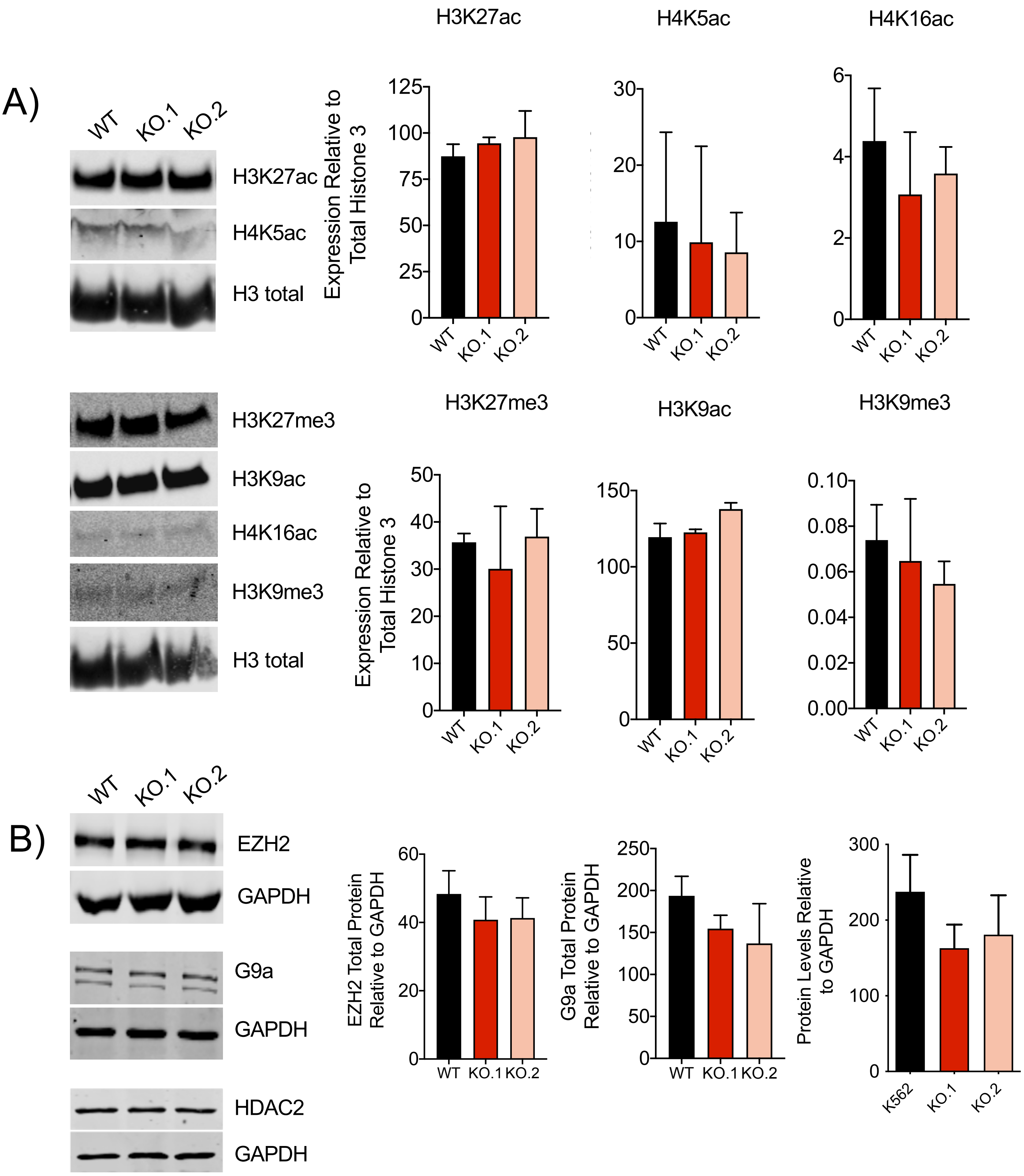


**Supplementary Figure 2:**

(A) Immunoblotting on histone protein isolated from WT and CCDC26 KO cells using antibodies against common histone modifications (anti-H3K27ac, -H4K5ac, -H3K27me3, -H3K9ac, -H4K16ac and H3K9me3). Levels were measured relative to total histone H3 levels. Values represent the mean ± standard deviation (n=3) (unpaired, two-tailed *t* test).

(B) Total protein levels of histone modifying enzymes, EZH2, G9a and HDAC2 measured relative to GAPDH by immunoblotting, are unchanged in WT and *CCDC26* KO cells. Values represent the mean ± standard deviation (n=3) (unpaired, two-tailed *t* test).

### SUPPLEMENTARY FIGURE -3

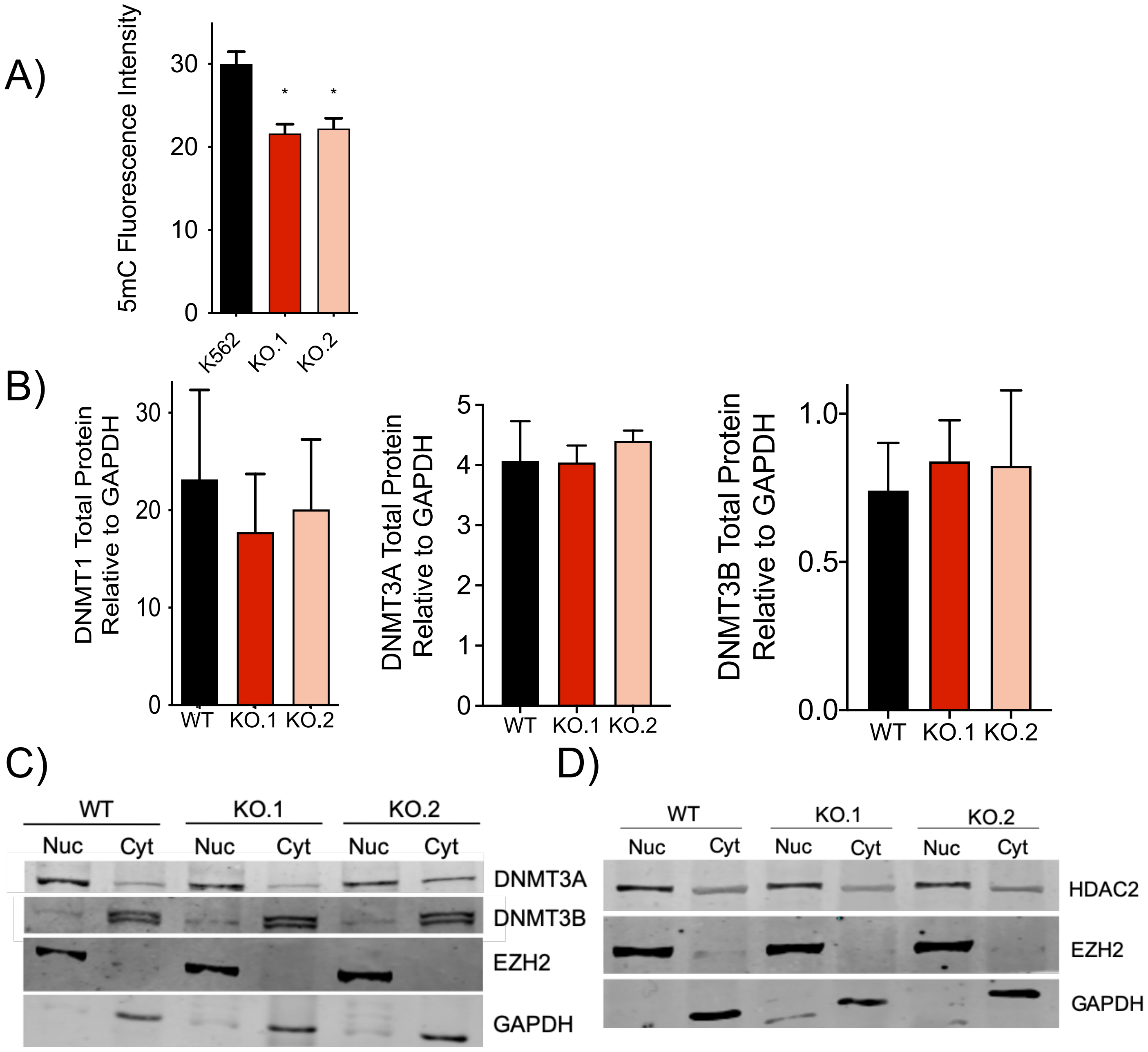


**Supplementary Figure 3:**

(A) 5mC Immunofluorescence intensity measurements in *CCDC26* KO cells compared to WT. 5mC fluorescence intensity was measured in 2D confocal images for 200 individual nuclei, per replicate, using FIJI image analysis software. Values represent the mean ± standard deviation (n=3). * P<0.05 (unpaired, two-tailed *t* test).

(B) Plots showing total protein levels of DNMT1, DNMT3a and DNMT3b relative to GAPDH in WT K562, KO.1 and KO.2 cells by immunoblotting. Values represent the mean ± standard deviation. * P<0.05 (unpaired, two- tailed *t* test) (n=3).

(C) Immunoblotting for DNMT3A and DNMT3B on nuclear and cytosolic protein fractions show no significant difference in subcellular localisation between WT and *CCDC26* KO cells. EZH2 and GAPDH are used as nuclear and cytosolic markers respectively (nuc = nuclear protein fraction; cyt = cytosolic protein fraction).

(D) Immunoblotting for HDAC2 on nuclear and cytosolic protein fractions show no significant difference in subcellular localisation between WT and *CCDC26* KO cells. EZH2 and GAPDH are used as nuclear and cytosolic markers respectively (nuc = nuclear protein fraction; cyt = cytosolic protein fraction).

### SUPPLEMENTARY FIGURE -4

A)

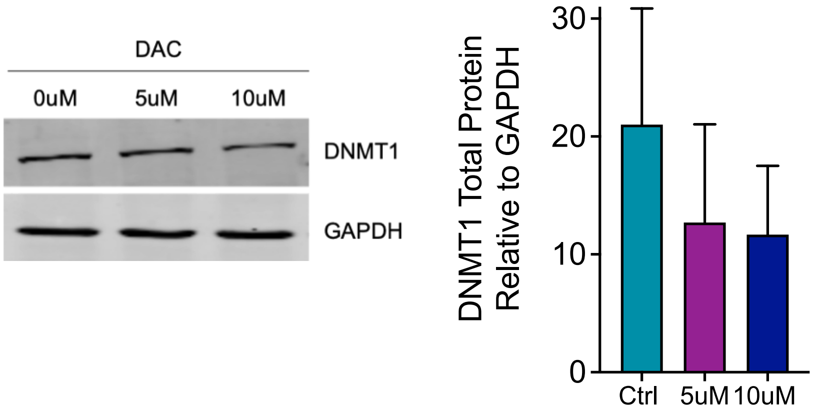


B)

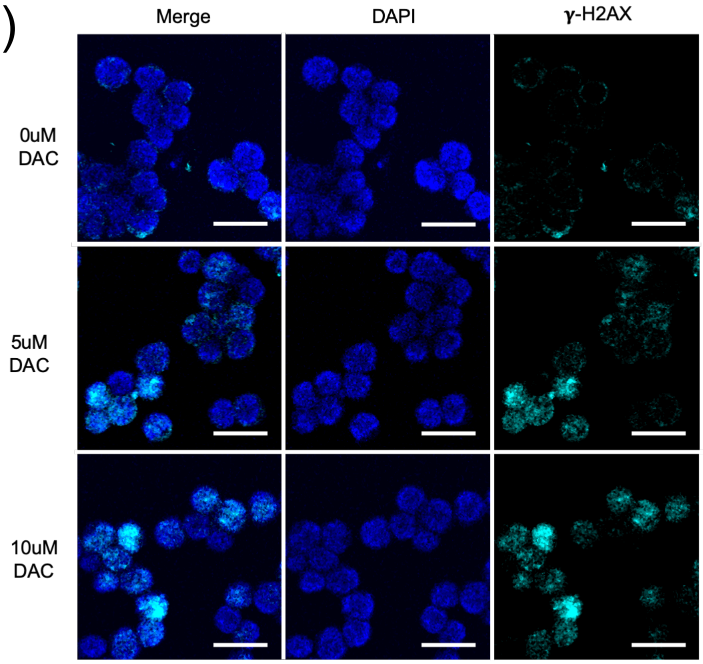


**Supplementary Figure 4:**

(A) Immunoblotting for DNMT1 total protein levels in cells treated with 0uM 5uM and 10uM DAC, measured relative to GAPDH. DNMT1 levels are slightly reduced in cells treated with 5uM and 10uM DAC. Values represent the mean ± standard deviation (n=3) (unpaired, two-tailed *t* test).

(B) Confocal images demonstrating the results of anti-**γ**-H2AX immunofluorescence. Cells treated with 0uM, 5uM and 10uM DNMT1 inhibitor, DAC, were stained with DAPI nuclear stain (blue) and anti-**γ**-H2AX antibody (cyan). Increased numbers of **γ**-H2AX foci are present in the cells treated with 5uM and 10uM DAC Scale bar = 25um.

### SUPPLEMENTARY FIGURE -5

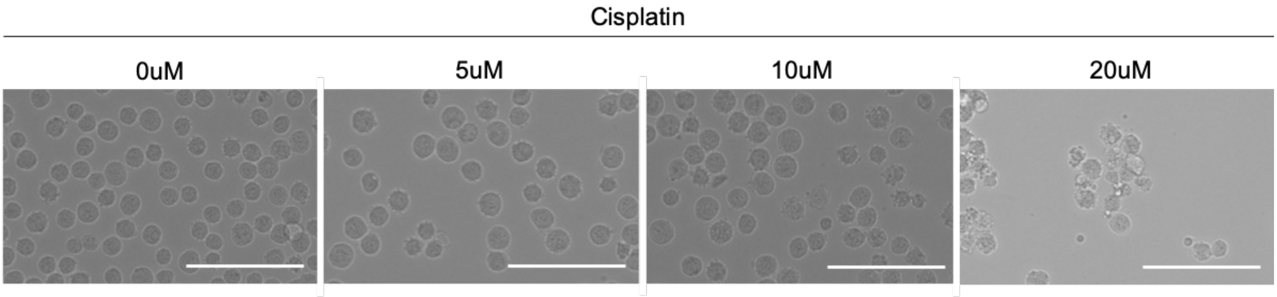


**Supplementary Figure 5:**

Brightfield microscopy images of K562 cells treated with increasing concentrations of DNA damage-inducing drug, cisplatin. Cells appear increasingly distressed with increasing cisplatin concentrations. Scale bar = 100um.

### SUPPLEMENTARY FIGURE -6

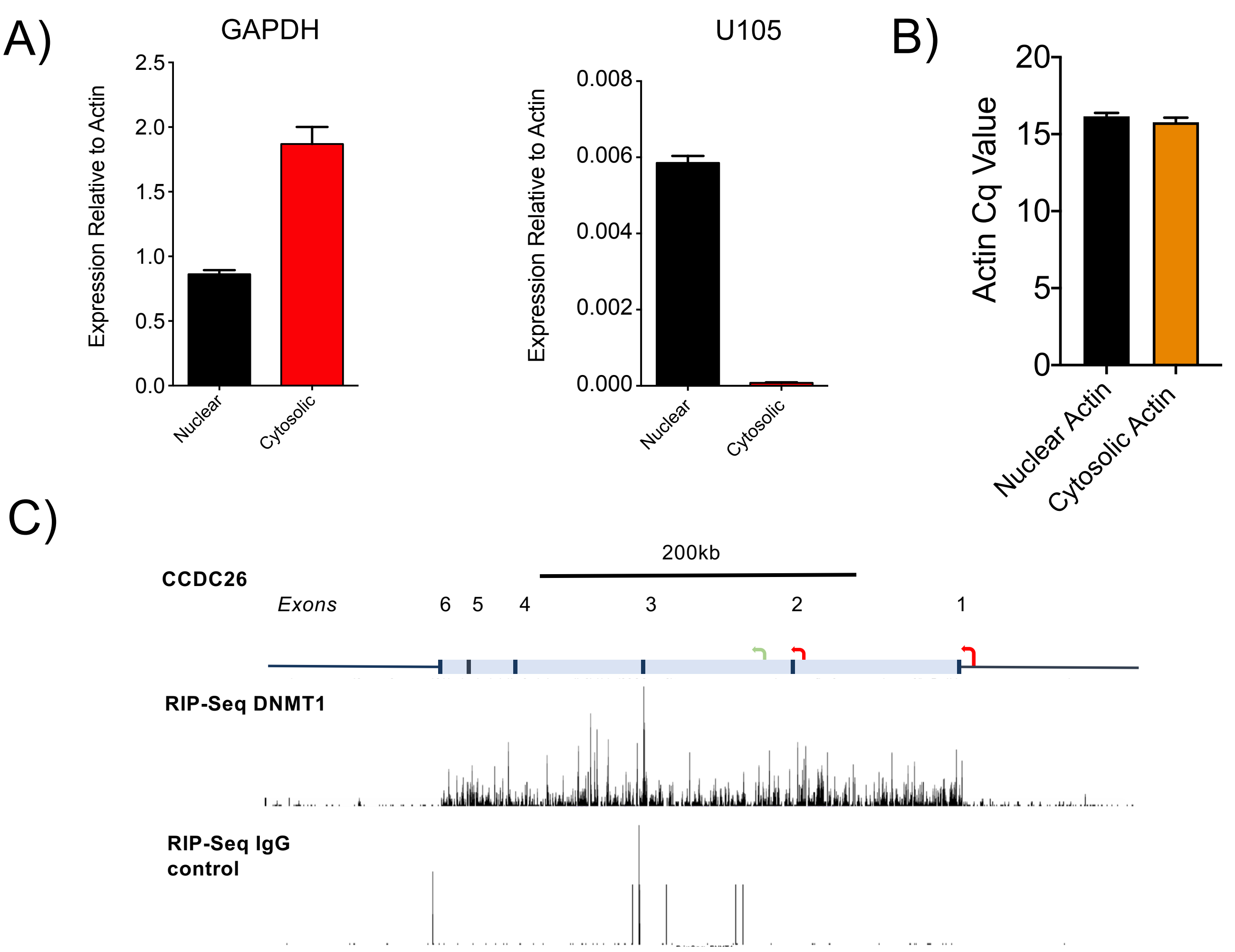


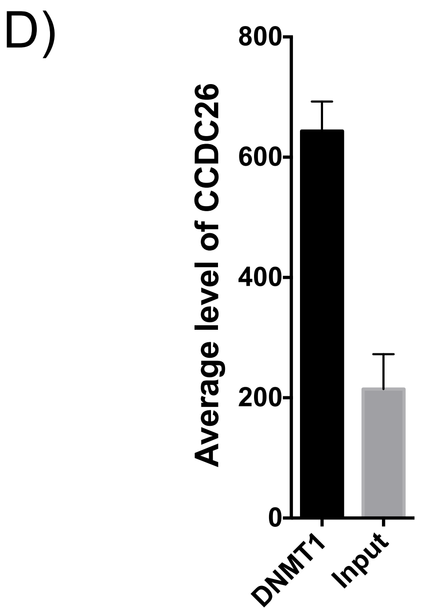


**Supplementary Figure 6:**

(A) Plots showing RNA levels of GAPDH and U105 in cytosolic and nuclear fractions relative to Actin level. Values represent the mean ± standard deviation. * P<0.05 (unpaired, two- tailed *t* test, n=3).

(B) Cq measurements for K562 nuclear and cytosolic RNA fractions generated by qRT-PCR using Actin primers.

(C) Re-analysis of a previously published DNMT1 RIP-seq data set, performed in HL60 cells. Re-mapping the cellular RNAs pulled down with either a DNMT1 antibody or an IgG control antibody shows enrichment for *CCDC26* in DNMT1-RNA, compared to IgG control. (Di Ruscio et al. 2013) (GEO Accession: GSE32162). Data available at NCBI Gene Expression Omnibus (GEO).

(D) Re-analysis of a previously published DNMT1 fRIP-seq data set, performed in K562 cells shows high enrichment of *CCDC26* in DNMT1-bound RNAs (Hendrickson et al. 2016) (GEO Accession: GSE67963). Data available at NCBI Gene Expression Omnibus (GEO).

### SUPPLEMENTARY FIGURE -7

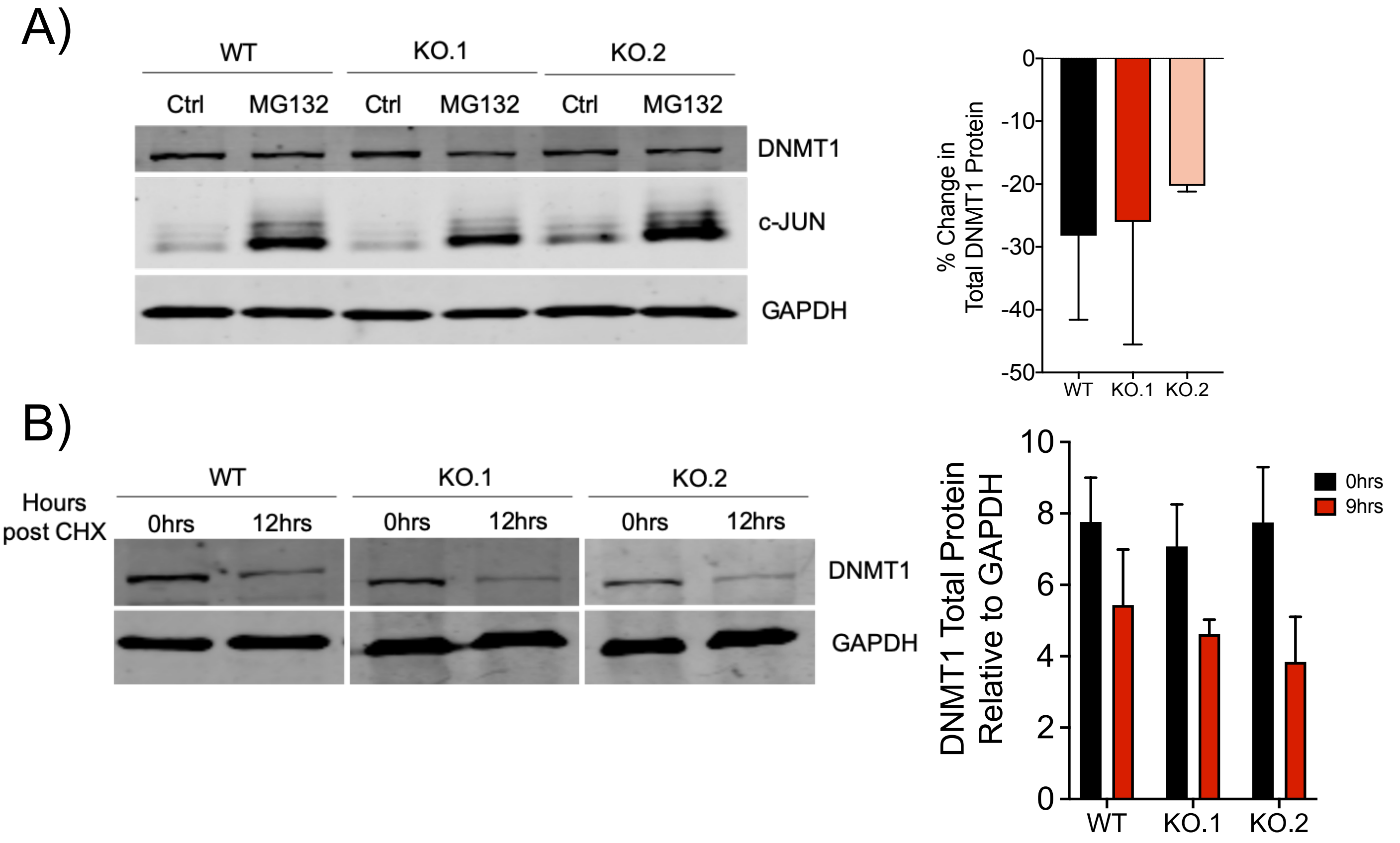


**Supplementary Figure 7:**

(A) Immunoblotting for DNMT1 total protein levels in WT and CCDC26 KO cells following treatment with 10uM of proteosomal inhibitor MG132. DNMT1 protein levels fell in both WT and KO cells. The difference in the extent to which DNMT1 levels fall was not statistically significant between cell lines. Immunoblotting for c-JUN was also performed as a control to show that the MG132 inhibitor was working. C-JUN levels rose upon MG132 treatment in both WT and KO cells. Protein levels were measured relative to the housekeeping protein, GAPDH. Values represent the mean ± standard deviation (unpaired, two-tailed *t* test, n=3).

(B) Immunoblotting for DNMT1 total protein levels on WT and CCDC26 KO cells treated with cycloheximide (CHX) for 0hrs and 12hrs. The difference in the extent to which DNMT1 levels fall was not statistically significant between WT and KO cells. Values represent the mean ± standard deviation (unpaired, two-tailed *t* test, n=3).
